# Lantibiotic-producing bacteria impact microbiome resilience and colonization resistance

**DOI:** 10.1101/2025.05.06.652565

**Authors:** Cody G. Cole, Zhenrun J. Zhang, Shravan R. Dommaraju, Qiwen Dong, Rosemary L. Pope, Sophie S. Son, Emma J. McSpadden, Che K. Woodson, Huaiying Lin, Nicholas P. Dylla, Ashley M. Sidebottom, Anitha Sundararajan, Douglas A. Mitchell, Eric G. Pamer

## Abstract

A subset of commensal bacterial strains secrete bacteriocins, such as lantibiotics, to establish and protect their niche in the gut. Because the antimicrobial spectrum of lantibiotics includes opportunistic pathogens, such as vancomycin-resistant *Enterococcus faecium* (VRE), they may provide an approach to reduce antibiotic-resistant infections. The impact of lantibiotic-producing bacteria on the complex microbial populations constituting the microbiome, however, remains poorly defined. We find that genes encoding lanthipeptides, including lantibiotics, are commonly present in the microbiomes of healthy humans and in dysbiotic microbiomes of hospitalized patients. In fecal samples collected from hospitalized patients, bacterial species encoding lantibiotic genes are present in greater abundance than lantibiotic-deficient strains of the same species. We demonstrate that the lantibiotic-producing bacterium, *Blautia pseudococcoides* SCSK, prevents intestinal recolonization of mice by a wide range of commensal species following antibiotic-induced dysbiosis and markedly reduces fecal concentrations of microbiota-derived metabolites associated with mucosal immune defenses. Lantibiotic-mediated dysbiosis results in sustained loss of colonization resistance against *Klebsiella pneumoniae* and *Clostrioides difficile* infection. Our findings reveal the potential impact of lantibiotic-producing bacterial species on microbiome resilience and susceptibility to infection following antibiotic treatment.

## Introduction

The mammalian gut microbiota is home to trillions of bacteria that, together, form a complex community with a vast array of microbe-microbe and host-microbe interactions. These dense interactions lead to competition between microbes for limited nutrients. To gain a competitive edge within their niche, some bacterial strains encode genes expressing narrow- and broad-spectrum bacteriocins that inhibit or kill neighboring bacteria by targeting specific cellular components or disrupting their bacterial cell membrane/wall^1^. Bacteriocins include non-ribosomal peptides (NRPs), ribosomally synthesized and post-translationally-modified peptides (RiPPs)^2,3^, and proteins. Metagenomic analysis has revealed that many gut-associated commensals encode biosynthetic gene clusters (BGCs) to produce a wide range of bacteriocins^4–8^.

Given their antimicrobial properties, bacteriocins such as nisin and bacitracin have been used extensively as food preservatives and topical treatments, respectively, to reduce the risk of bacterial infections^9,10^. With the rise of antibiotic resistance among human pathogens, there is growing interest in developing bacteriocins as new antibiotic agents, either as purified drugs or as bacteriocin-producing probiotics^11–16^. Therapeutic bacteriocins might be administered to clear pathogens during active infection or used preemptively to prevent infection by pathogens in vulnerable patients.

While progress has been made in identifying bacteriocins with activity against pathogens, the extent to which bacteriocin-producing bacteria shape and modulate the gut microbiota remains unclear. Some studies have explored the impact of bacteriocins–particularly lantibiotics–on the microbiota compositions or specific microbes^17–25^. Given the broad activity of bacteriocins, their expression likely influences microbiota composition, particularly during recovery from states of dysbiosis, by preventing recolonization with bacteriocin-sensitive commensals. Additionally, if the species that cooperate and support the growth of others in the community are prevented from recolonizing, it might, in turn, indirectly prevent colonization by gut commensals dependent on those species.

Lantibiotics are a subset of lanthipeptides that have antimicrobial properties. Lanthipeptide BGCs represent one of the most prevalent BGCs in the human microbiome^6,8,26^. Currently, there are six classes of lanthipeptides that are classified based on the biosynthesis genes required to produce the lanthipeptide^2,27,28^. Class I lanthipeptides have two synthesis enzymes that consist of LanB, which dehydrates serine and threonine residues, and LanC, which cyclizes the *β*-carbon of dehydroamino acids with cystine to form the characteristic lanthionine and/or 3-methyllanthionine bonds. Class II, III, IV, and VI use single multifunctional synthetases to carry out the same processes. These proteins are LanM (class II), LanKC (class III), LanL (class IV), and LanKCb (class VI). Class V uses the enzymes LanK, LanY, and LanC. Although lanthipeptides can serve multiple functions, many have antimicrobial activity, especially class I and II lanthipeptides^3^.

Herein, we demonstrate that lanthipeptide-encoding genes are common in fecal metagenomes of healthy humans and hospitalized patients with varying degrees of dysbiosis. In comparison to bacterial strains lacking lanthipeptide-encoding genes, those encoding them are more abundant, suggesting they contribute to fitness in the gut. Lanthipeptide genes were detected in several species including *Blautia* species and probiotic strains such as *Streptococcus thermophilus*, *Bifidobacterium longum*, *Bifidobacterium breve*, *Lactiplantabacillus plantarum*, and *Lactococcus lactis*, which have been shown to prolong dysbiosis after antibiotic treatment by an undefined mechanism^29^. Additionally, we show that *Blautia pseudococcoides* SCSK (BpSCSK), a lantibiotic-producing bacterium that provides colonization resistance against vancomycin-resistant *Enterococcus faecium* (VRE), prolongs antibiotic-induced dysbiosis by preventing recolonization of the gut microbiota with gut commensals^30,31^ and reduces fecal metabolite concentrations associated with epithelial barrier function and mucosal immune defenses. In antibiotic-treated mice, BpSCSK prolongs susceptibility to gut colonization with *Klebsiella pneumoniae* and infection with *Clostrioides difficile*. These findings highlight the potentially detrimental role bacteriocin-producing commensal bacteria can play in microbiota diversification and function.

## Results

### Lanthipeptide gene prevalence and diversity in patient microbiomes

Patients hospitalized for respiratory failure, sepsis, or liver disease/transplantation are frequently treated with broad-spectrum antibiotics that disrupt the gut microbiota. Upon completion of antibiotic treatment, commensal bacterial species recolonize the gut at variable rates that are likely determined by the patient’s environmental exposures and the composition of the residual microbiota. During the reconstitution phase, expression of lantibiotics by some strains might enhance their fitness in the gut lumen. We collected fecal samples from patients undergoing hospitalization for critical care, organ transplantation, and liver disease, and subjected them to shotgun metagenomic sequencing and metabolomic analyses, as previously described^32–34^. Using a combination of RODEO (Rapid ORF Description and Evaluation Online), previously identified lanthipeptide gene sequences, and specific lanthipeptide-associated Pfam domains^7,35^, we screened 1872 sequenced fecal samples from 612 patients and 17 healthy donors for the presence of lanthipeptide encoding genes.

Fecal microbiomes from 526 out of 612 patients (∼86%) contained at least one detectable lanthipeptide precursor encoding gene (*lanA*) **(Fig. 1a)**. Of the 1872 fecal samples screened, 71% contained at least one *lanA* gene, yielding 1015 unique lanthipeptide sequences. Lanthipeptide classes were determined on the basis of gene annotations or neighboring synthesis genes. Although classes of some lanthipeptides could not be determined, most of the *lanA* sequences detected belonged to class II lanthipeptides (640/1015) followed by class I (231/1015) and III (36/1015) **(Fig. 1b)**. All healthy donors contained at least one *lanA* sequence with 59 unique sequences detected in 17 healthy donor microbiomes.

**Fig. 1.**
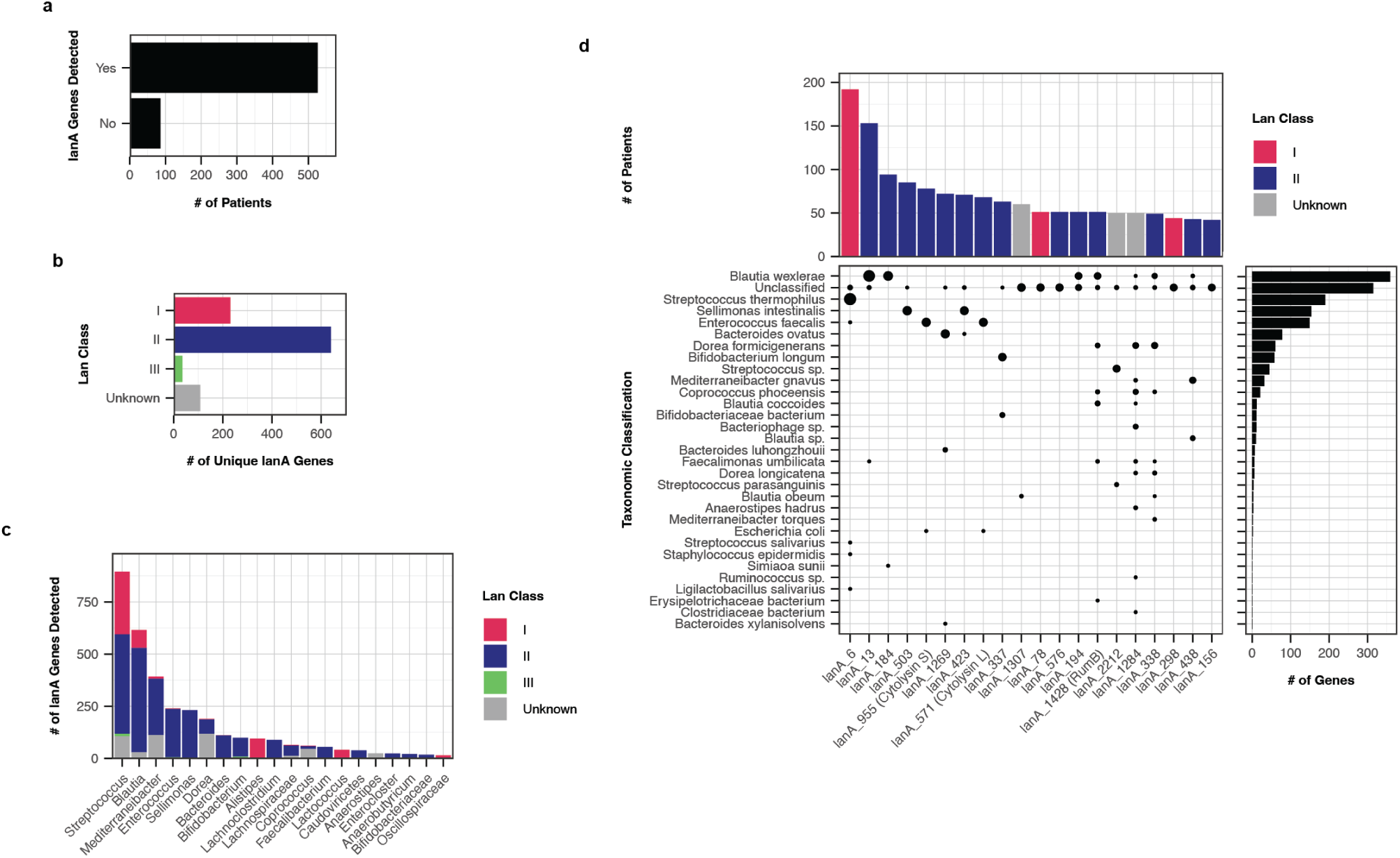
Lanthipeptide gene prevalence and diversity in patient microbiomes. A total of 1872 fecal samples were collected from 612 patients and screened for lanthipeptide genes. **a,** Patients that had a detectable lanthipeptide gene among fecal samples sequences. **b,** Lanthipeptide class distribution for unique lanthipeptides detected in patient samples (*n* = 1015). **c,** Distribution of detected lanthipeptide genes across the top 20 genera assigned to the encoding contig. **d,** Plot of the top 20 most abundant lanthipeptide genes. Vertical columns represent the number of times the lanthipeptide was detected. Horizontal bars represent the number of lanthipeptides were found in a given species. The dots represent the intersection between the two plots with the size of the dots representing the number of times a specific lanthipeptide was found in a specific species. Previously published lanthipeptide core sequences that were a match to the identified lanthipeptide are specified in parentheses.

Taxonomic classifications for *lanA* sequences were determined using NCBI BLAST on contigs harboring these genes. Contigs that did not meet the inclusion thresholds or did not have any hits were labeled “Unclassified.” Out of 5503 contigs, 3869 contigs were assigned taxonomic classification. At the genus level, *Streptococcus*, *Blautia*, and *Mediterraneibacter* were the most prominent genera **(Fig. 1c)**. Most *lanA* sequences were predominantly detected in one species while some, such as lanA 1428 (RumB), lanA 1284, and lanA 338, were distributed across multiple species, suggesting they may have been acquired by horizontal gene transfer **(Fig. 1d)**. At the species level, *Blautia wexlerae* contained the most *lanA* sequences, followed by *Mediterraneibacter gnavus* and *Streptococcus thermophilus* **(Extended Data Fig. 1a)**.

While *B. wexlerae* and *M. gnavus* are common human gut commensals, *S. thermophilus* is typically associated with fermented milk and is used in yogurt production. Other species with *lanA* genes that are commonly used as probiotics included *Bifidobacterium longum*, *Lactiplantabacillus plantarum*, *Lactococcus lactis*, *Ligilactobacillus salivarius*, and *Bifidobacterium breve*. Lanthipeptides were also detected in oral microbiome associated *Streptococcus salivarious*, *Streptococcous parasanguinis*, *Streptococcos mutans*, and *Streptococcous pneumoniae* in addition to the nasal mucosal and dermal microbiome associated *Staphylococcus aureus* and *Staphylococcus epidermidis* **(Extended Data Fig. 1a)**.

Whether bacterial strains encoding *lanA* genes impact microbiota composition, however, is unclear, since only a subset of lanthipeptides have been demonstrated to have antibacterial activity and some encoded genes may not be transcribed in the gut. To determine whether lantibiotics provide a fitness advantage to bacterial species expressing them, we assessed the impact of the *lanA* genes in patient microbiomes by calculating the ratio of the average relative abundance of species with a *lanA* gene to the average relative abundance of the same species lacking the *lanA* gene. Since not all *lanA* contig taxonomic classifications could be matched to a species annotated by MetaPhlAn4^36^ within a sequenced sample, only exact matches at the species level were considered. Moreover, only non-producing species and *lanA* genes that were present in at least 5 samples were used. Among the 1015 unique lanthipeptide sequences in patients, 102 sequences met the criteria for inclusion. We found that 80 out of the 102 lanthipeptide encoding strains had increased abundance compared to lanthipeptide lacking strains **(Fig. 2a)**. A UMAP of all the lanthipeptide sequences revealed that there was no clear clustering of lanthipeptides that provided a fitness advantage **(Fig. 2b)**. However, class I and class II lanthipeptides largely separated along the axis 2 of the UMAP.

**Fig. 2.**
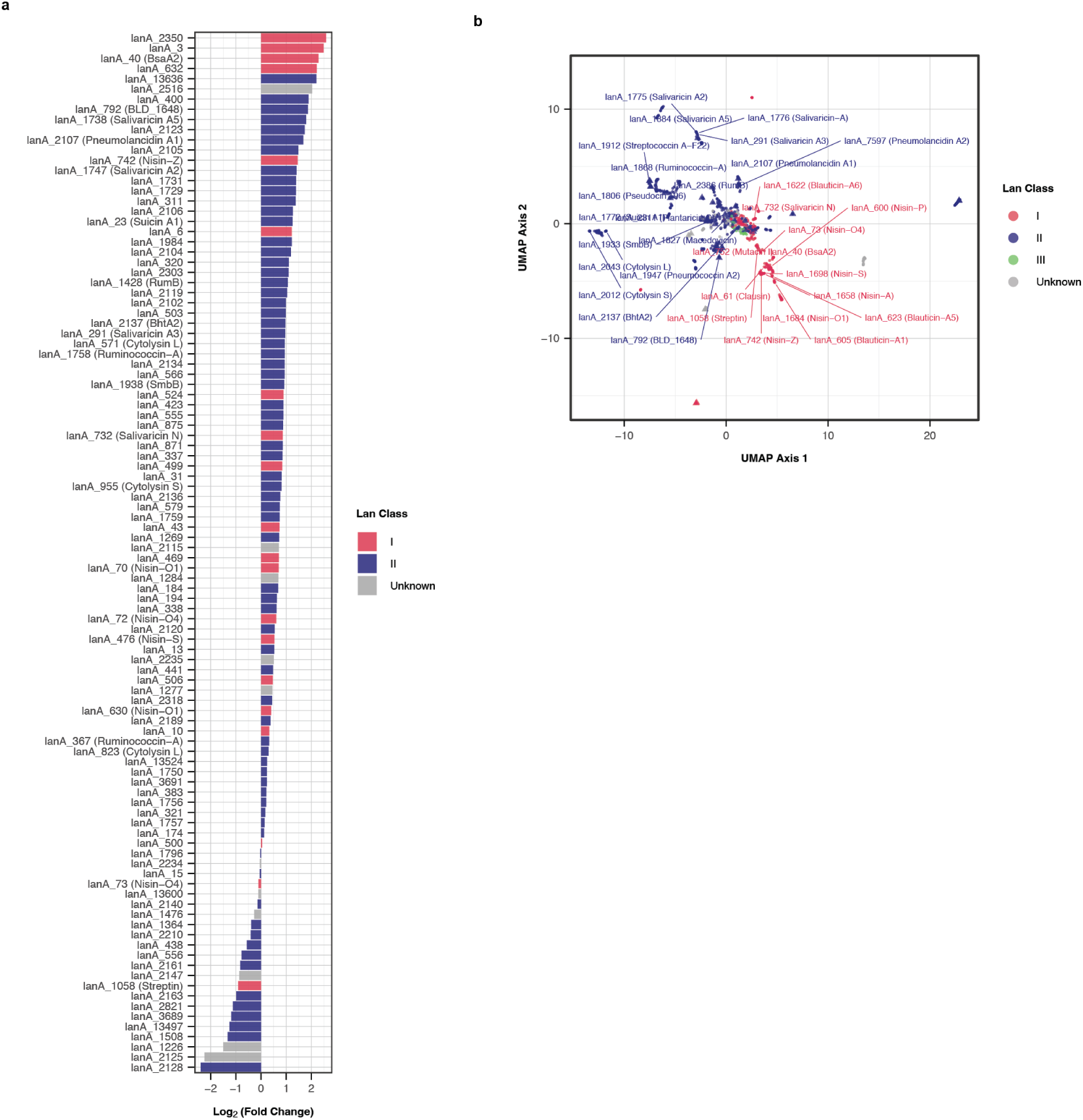
Lanthipeptides provide a fitness advantage for producing bacteria. **a,** Fitness advantage of lanthipeptide genes calculated as the log_2_ fold change of the relative abundance of species with the lanthipeptide gene versus the relative abundance of the same species without the lanthipeptide gene. Only species that were found in at least 5 fecal samples with and without the lanthipeptide gene were included. **b,** UMAP of all unique lanthipeptide genes detected across all samples. Triangles indicate lanthipeptide genes with a log_2_ fold change greater than 0. Previously published lanthipeptide core sequences that were a match to the identified lanthipeptide are specified in parentheses.

Among the 1015 lanthipeptide sequences, 105 matched the core sequences of 33 previously characterized lanthipeptides. The most abundant were the two cytolysin L and S lanthipeptides associated with *Enterococcus faecalis* **(Extended Data Fig. 1b)**. Unexpectedly, three of the lanthipeptides detected, lanA 605 (Blauticin-A1), lanA 623 (Blauticin-A5), and lanA 1622 (Blauticin-A6), were exact matches to the class I lantibiotic genes of *Blautia pseudococcoides* SCSK (BpSCSK). Previous research has shown that blauticin from BpSCSK, an anaerobic, Gram-positive, spore-forming strain of *Blautia* in the *Lachnospiraceae* family, inhibits VRE^30,31^. To determine the impact of commensal bacterial expression of an active lantibiotic on microbiome composition and function, we focused on the impact BpSCSK colonization has on the gut microbiome and metabolome.

### BpSCSK suppresses microbiota recolonization following antibiotic treatment

Lantibiotic expression by BpSCSK promotes VRE clearance from the gut^30,31^, but the extent to which it suppresses commensal bacterial populations and potentially enables BpSCSK to invade a diverse gut microbiota is unclear. To address this, we administered BpSCSK by oral gavage daily for three days to specific pathogen-free (SPF) C57BL/6 mice from Jackson Laboratories **(Extended Data Fig. 2a)**. Since genetic tools to manipulate BpSCSK are currently lacking, we gavaged mice with the closely related strain *Blautia producta* KH6 (BpKH6) strain, which does not encode a lantibiotic biosynthetic gene cluster (BGC), as a negative control. Neither BpSCSK nor BpKH6 were detected in fecal samples by 16S rRNA gene sequencing in recipient mice harboring the SPF microbiota **(Extended Data Fig. 2b)**. Thus, BpSCSK’s ability to express blauticin did not enable it to invade and colonize a diverse SPF microbiota following repeated oral gavage.

To reduce microbiota-mediated colonization resistance, mice were treated with 0.5 g/L ampicillin in drinking water for four days to deplete the gut microbiota. Two days after ampicillin treatment was stopped, mice were orally gavaged with BpSCSK, BpKH6, or a PBS vehicle control for three consecutive days **(Fig. 3a)**. Following oral gavage, both BpSCSK and BpKH6 colonized the gut post-ampicillin treatment and persisted for up to 28 days **(Fig. 3b)**. BpSCSK represented *>*50% of the 16s rRNA gene relative abundance of fecal pellets collected on all days following oral gavage, while BpKH6 had a lower abundance upon colonization and continued to decrease overtime. Unlike the PBS- and BpKH6-treated groups, all mice colonized with BpSCSK maintained a significantly lower *α*-diversity for the 28 days mice were monitored, as evidenced by the reduced unique amplicon sequence variants (ASVs) and lower Shannon diversity index **(Fig. 3c and 3d)**. PBS- and BpKH6-treated mice reached similar levels of *α*-diversity after 1-3 days post-colonization; however, both remained slightly lower than those of untreated, antibiotic-naive SPF mice after 28 days. Furthermore, PBS- and BpKH6-treated groups clustered more closely with SPF mice than BpSCSK-colonized mice based on their Bray-Curtis dissimilarity **(Fig. 3e)**. Despite the differences observed in diversity, all mice had similar overall bacterial densities in feces, as determined by RT-qPCR **(Extended Data Fig. 3a)**.

**Fig. 3.**
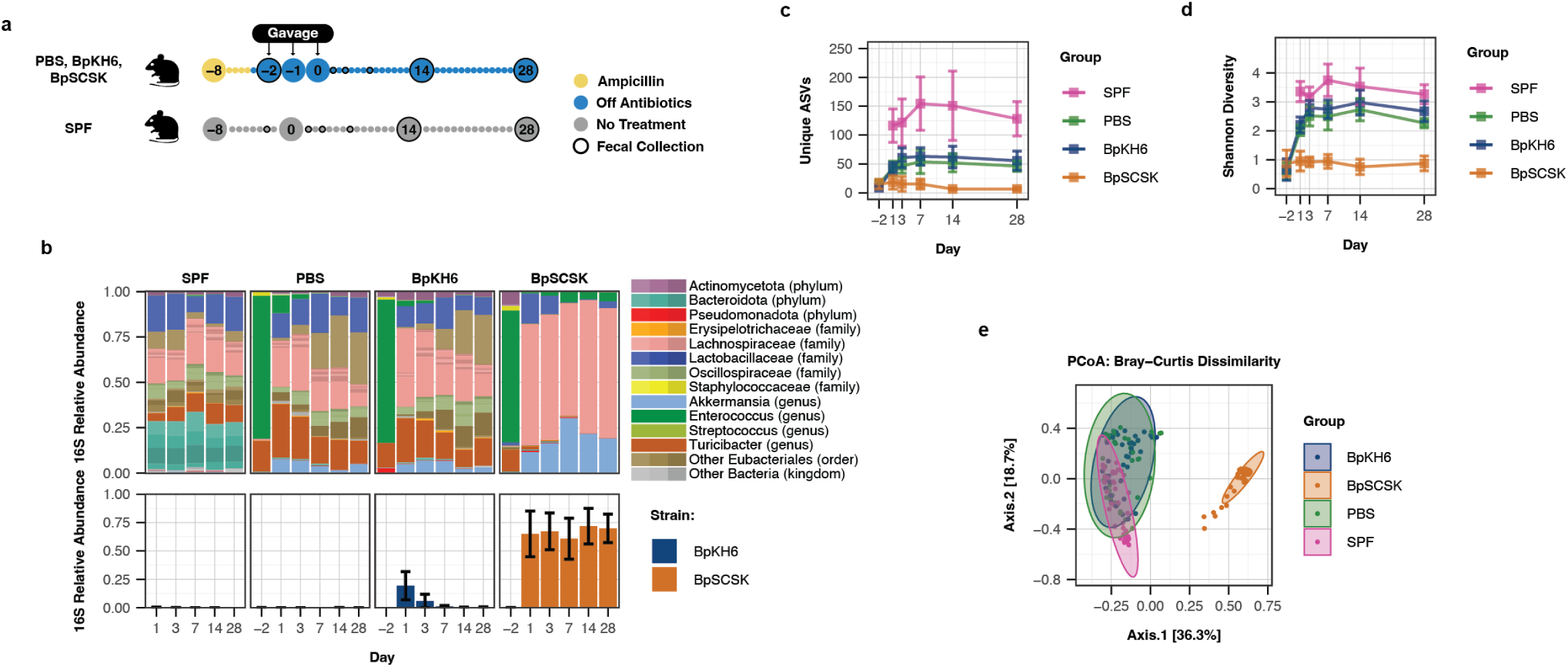
BpSCSK suppresses microbiota recolonization following antibiotic treatment. **a,** Schematic of experimental groups and time points. Female wild-type SPF C57BL/6 mice 9-10 weeks old were treated with 0.5 g/L ampicillin in their drinking water for 4 days and given PBS, BpKH6, or BpSCSK via oral gavage. Fecal samples were collected at indicated time points. Each group contains 8 mice. **b,** Average fecal microbiota 16S rRNA gene relative abundance. ASVs with greater than *>*0.01% abundance are plotted. Specific ASVs for BpSCSK (orange) and BpKH6 (blue) are plotted below. Error bars indicate standard deviation. **c,** Average number of unique ASVs detected. Error bars indicate 95% confidence intervals. **d,** Average Shannon diversity. Error bars indicate 95% confidence intervals. **e,** Bray-Curtis dissimilarity principal coordinates analysis (PCoA) for all time points excluding day −2.

Following ampicillin treatment, *Enterococcus* and *Turicibacter* were among the most abundant genera detected in fecal pellets **(Fig. 3b)**. In BpSCSK-colonized mice, *Turicibacter* diminished over the 28-day period while the *Enterococcus* and *Lactobacillus* ASVs persisted along with *Akkermansia*, which was detected from day 1 post gavage. Among ASVs across all groups, ASVs that belong to the Eubacteriales order and particularly within the *Lachnospiraceae* and *Oscillospiraceae* families represented the majority of ASVs that did not recolonize mice containing BpSCSK after 28 days, including the ASVs for *Lachnospiraceae* and *Oscillospiraceae* that were present prior to the introduction of BpSCSK **(Fig. 3b**; **Table 1)**. Of the *Lachnospiraceae* ASVs that remained in BpSCSK-colonized mice after 28 days, all 10 belonged to the *Blautia* genus. These findings demonstrate that the lantibiotic-producing BpSCSK strain markedly impairs recolonization by gut commensals following antibiotic treatment.

**Table 1.**
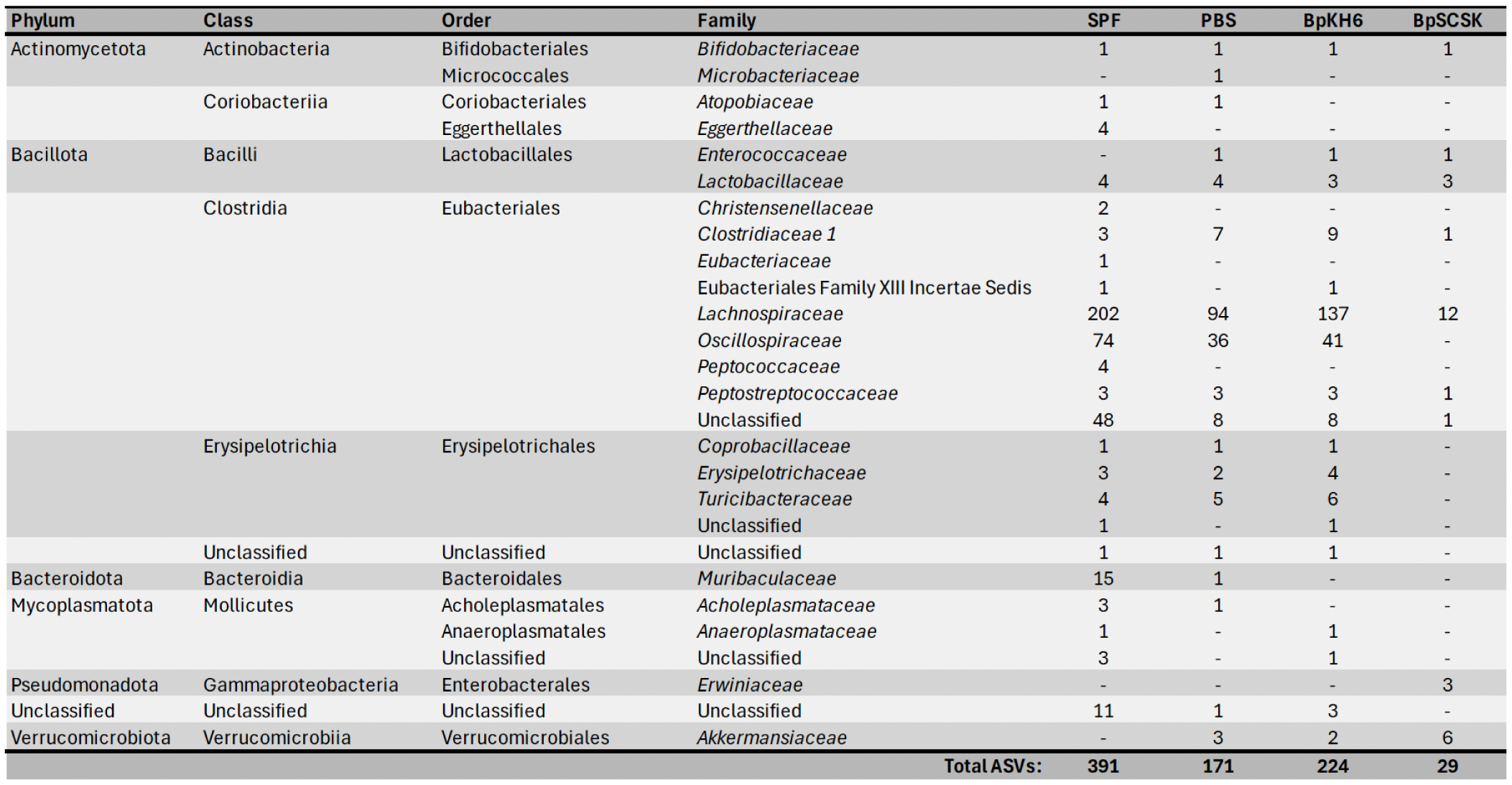
Unique ASVs detected on day 28 of colonization. Taxonomic distribution of ASVs from 16S rRNA gene sequencing of mice at day 28 in Fig. 3. Values indicate the number of unique ASVs at the family level detected at *>*0.01% relative abundance.

### The fecal metabolome is rendered deficient by BpSCSK colonization

*Lachnospiraceae* and *Oscillospiraceae* encode genes that enable production of microbiota-derived metabolites, including short-chain fatty acids (SCFAs) and a range of bile acid variants^37^. Since the species from these families were not well represented in the fecal pellets of mice harboring BpSCSK compared to the PBS and BpKH6 controls, we investigated the metabolomic profiles of fecal pellets collected over 28 days from these mice.

Using gas chromatography-mass spectrometry (GC-MS) with pentafluorobenzyl-bromide (PFBBr) derivatization, we determined the concentrations of a panel of 8 compounds that include amino acids, fatty acids, and organic acids. As expected, 4-day treatment with ampicillin led to the marked reduction of SCFA concentrations in all mouse groups compared to SPF mice **(Fig. 4a)**. In contrast, proline and tyramine concentrations increased after ampicillin treatment compared to SPF mice, while glycine concentrations only changed minimally. Within the first week following PBS- or BpKH6-treatment, concentrations of proline, butyrate, and acetate returned to those detected in SPF mice. Concentrations of 5-aminovalerate exceeded the relative concentrations in SPF mice by day 1 and continued to day 28, suggesting that antibiotic treatment resulted in the expansion of bacterial species that produce this metabolite. In contrast, mice colonized with BpSCSK had consistently lower concentrations of 5-aminovalerate and butyrate compared to SPF mice over the 28 days samples were collected. In addition, concentrations of glycine, proline, and succinate remained elevated for 28 days. The absolute concentrations of metabolites at day 28 revealed that BpSCSK-colonized mice did not have any detectable butyrate in their fecal pellets and confirmed the higher concentrations of glycine and proline **(Extended Data Table 1)**.

**Fig. 4.**
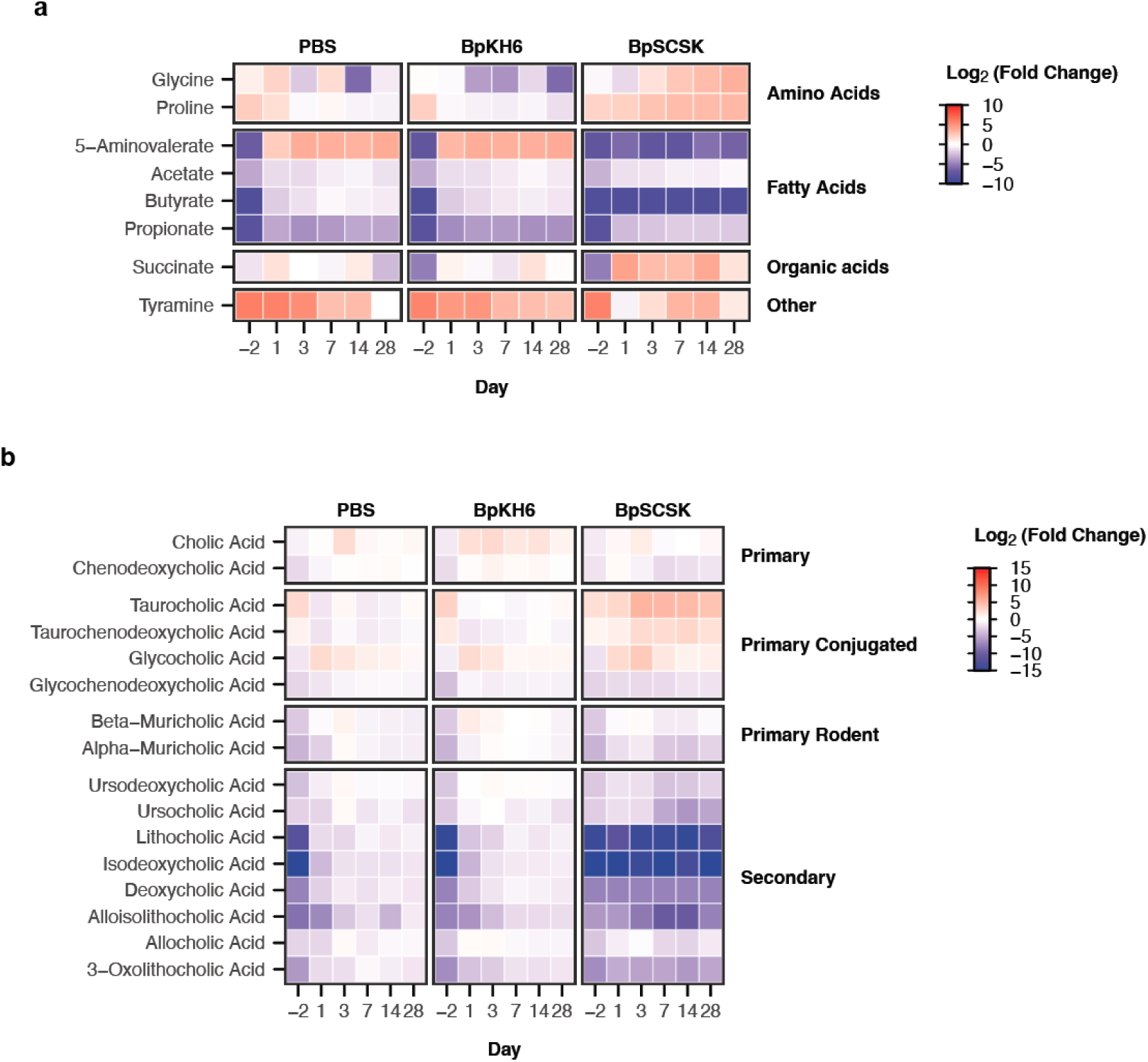
The fecal metabolome is rendered deficient by BpSCSK colonization. Fecal pellets collected from mice at the same time as 16S rRNA gene sequencing in Fig. 3 were subject to quantitative metabolomic profiling of a panel of metabolites. **a, b,** log_2_ fold change of metabolites calculated as the ratio of the average concentration of metabolites for indicated groups by the average concentration of metabolites across all days for SPF mice.

Next, we investigated the concentration changes of bile acids in fecal pellets using liquid chromatography-mass spectrometry (LC-MS). Before oral gavage, ampicillin treatment resulted in increased concentrations of conjugated primary bile acids, taurocholic and taurochenodeoxycholic acid, and decreased unconjugated primary bile acids compared to SPF mice **(Fig. 4b)**. In conjunction, the secondary bile acid concentrations began to decrease and secondary bile acids recovered and reached concentrations approaching those of SPF mice after termination of ampicillin treatment and were unaffected by BpKH6 or control PBS administration. In BpSCSK-colonized mice, in contrast, taurocholic and taurochenodeoxycholic acid levels remained elevated and secondary bile acid concentrations remained low up to 28 days after termination of ampicillin treatment. The relative abundance of *Turicibacter*, a genus with species previously shown to deconjugate taurine-conjugated primary bile acids, decreased during the course of BpSCSK colonization **(Fig. 3a)**^38^. Absolute concentrations of bile acids revealed that deoxycholic and isodeoxycholic acid were below the limits of detection, while other secondary bile acids were below the limits of detection, while other secondary bile acids were near zero except for ursodeoxycholic and allocholic acid **(Extended Data Table 1)**. Taurocholic acid concentrations were more than 10-fold greater in BpSCSK colonized mice than PBS- or BpKH6-treated mice.

Together, these metabolomic panels demonstrate that, in addition to decreased microbiota *α*-diversity, BpSCSK-colonized mice fail to recover critical producers of microbiota-derived metabolites. Therefore, BpSCSK prolongs both the compositional and functional dysbiotic state of mice up to 28 days after ampicillin treatment has ceased.

### Fecal microbiota transplantation fails to restore gut microbiota diversity and key metabolite producers

Recolonization of the gut microbiota following ampicillin treatment is driven by bacterial species that persist at low densities within the host or that are acquired from the surrounding environment. Therefore, it is possible that BpSCSK-colonized mice do not achieve the same level of diversity observed in PBS- and BpKH6-treated mice because they are unable to acquire commensal species from the environment. To address this, we attempted to recolonize mice by administering a fecal microbiota transplant (FMT) from antibiotic-naive SPF donor mice. Fecal pellets from four separate SPF mice were pooled and administered to BpSCSK-colonized mice via oral gavage for three consecutive days starting 28 days after the final gavage of PBS, BpKH6, or BpSCSK **(Fig. 5a)**. To demonstrate FMT viability, a separate group of mice were treated with ampicillin for four days starting six days before FMT **(Extended Data Fig. 4a)**.

**Fig. 5.**
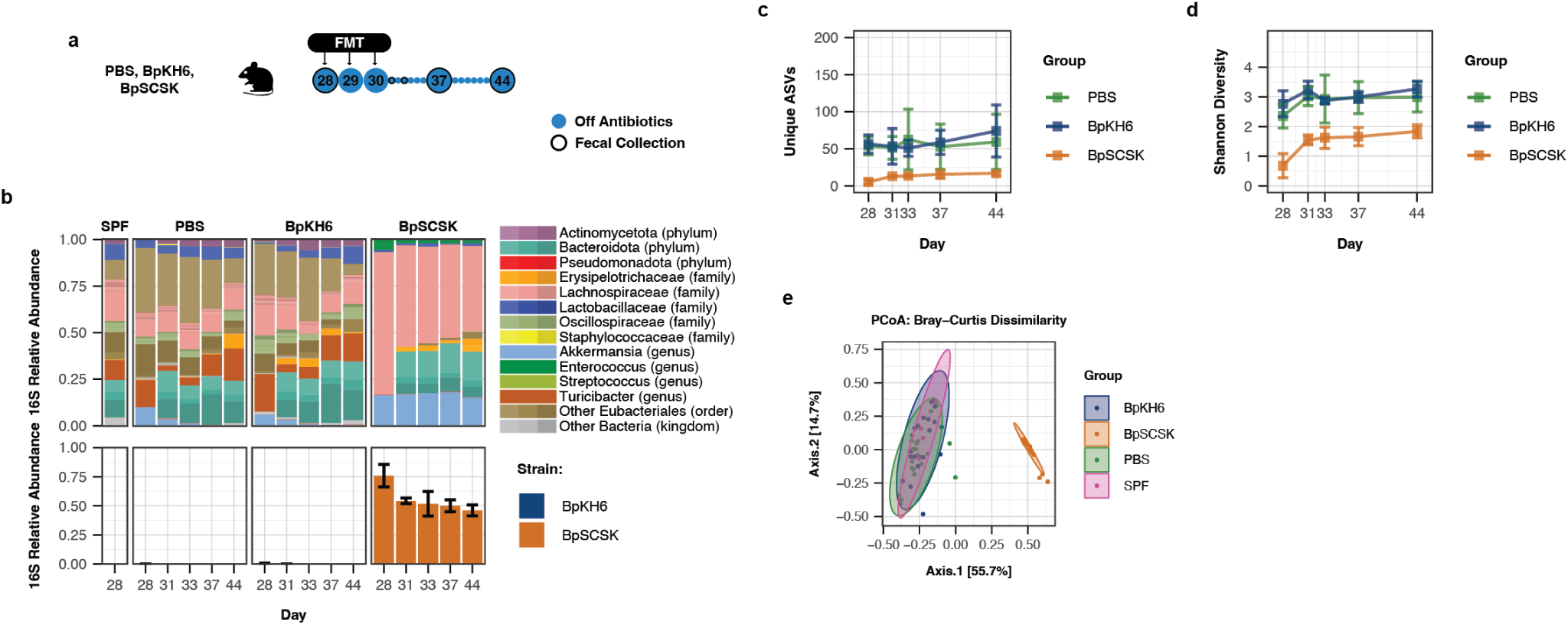
Fecal microbiota transplantation fails to restore gut microbiota diversity and key metabolite producers. **a,** Schematic of experimental groups and time points. 4 mice in each group from Fig. 3 were subjected to an FMT from an SPF mouse for three days via oral gavage. The FMT Control group were subject to 4 days ampicillin treatment starting 6 days before the first FMT. **b,** Average fecal microbiota 16S rRNA gene relative abundance. ASVs with greater than *>*0.01% abundance are plotted. Specific ASVs for BpSCSK (orange) and BpKH6 (blue) are plotted below. Error bars indicate standard deviation. **c,** Average number of unique ASVs detected. Error bars indicate 95% confidence intervals. **d,** Average Shannon diversity. Error bars indicate 95% confidence intervals. **e,** Bray-Curtis dissimilarity principal coordinates analysis (PCoA) for all time points.

In BpKH6-colonized mice, BpKH6 represented *<*0.5% of the relative abundance on day 28 and further decreased following FMT **(Fig. 3b and 5b)**. The PBS- and BpKH6-treated mice gained slightly more *α*-diversity after the FMT and reached a similar level of ASVs and Shannon diversity as FMT control mice **(Fig. 5c, 5d and Extended Data Fig. 4b)**. After three daily FMTs, BpSCSK still represented approximately 40% of the relative abundance of ASVs in fecal pellets after two weeks **(Fig. 5b)**. ASVs for *Muribaculaceae* and *Erysipelotrichaceae*, which were not detected before the FMT, made up approximately 35% in abundance on day 44. To a lesser extent, ASVs for *Oscillospiraceae* and an unknown Eubacteriales species were also detected on day 44. Although the Shannon diversity of BpSCSK-colonized mice was almost two-fold greater one day after the final FMT was administered, it remained at a Shannon diversity index of approximately 2 on day 44 while other groups had an average Shannon diversity index above 3 **(Fig. 5c and 5d)**. This was further highlighted by the Bray-Curtis dissimilarity between samples where the BpSCSK samples remained distinct from all other groups while PBS- and BpKH6-treated mice overlapped with samples from SPF and FMT control mice **(Fig. 5e)**.

Metabolomic profiling of mice before and after FMT treatment revealed that metabolites in PBS- and BpKH6-treated mice returned to the concentrations observed in SPF and FMT control **(Extended Data Fig. 5a and 5b)**. BpSCSK-colonized mice had increased concentrations of propionate, ursocholic acid, ursodeoxycholic acid, *α*-muricholic acid, *β*-muricholic acid, and chenodeoxycholic acid and decreased concentrations of taurocholic and taurochenodeoxycholic acid. Interestingly, the decrease in taurocholic and taurochenodeoxycholic acid was not associated with reestablishment of *Turicibacter* species **(Fig. 5b)**. Therefore, there are likely other bacteria among the *Muribaculaceae* and *Erysipelotrichaceae* deconjugate taurine. Other metabolites in the two panels were within similar ranges to the starting concentrations on day 28 in BpSCSK-colonized mice two weeks after FMT treatment **(Extended Data Fig. 5a and 5b)**. Thus, BpSCSK, once established in the gut, prevents colonization by many bacterial species following FMT from SPF mice.

### Co-housing BpSCSK-colonized mice with SPF mice fails to restore gut diversity and key metabolite producers

To determine whether a more regular introduction of bacteria from SPF mice via coprophagia could overcome colonization resistance mediated by BpSCSK, we co-housed BpSCSK-colonized mice with SPF mice. Each BpSCSK-colonized mouse was co-housed with one SPF mouse and fecal pellets were collected for 2 weeks for 16S rRNA gene sequencing and metabolomic profiling **(Fig. 6a)**.

**Fig. 6.**
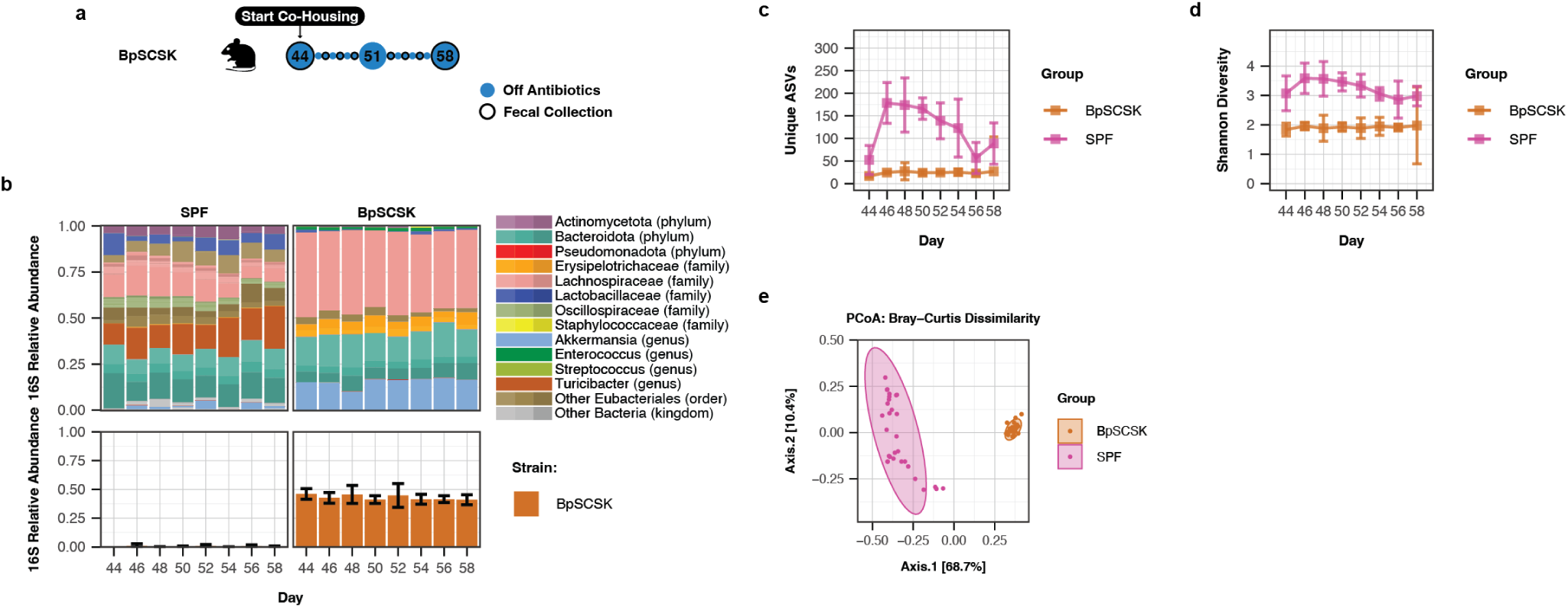
Co-housing BpSCSK-colonized mice with SPF mice fails to restore gut diversity and key metabolite producers. **a,** Schematic of experimental groups and time points. BpSCSK-treated mice from Fig. 5 that received an FMT were co-housed with an SPF mouse for two weeks. **b,** Average fecal microbiota 16S rRNA gene relative abundance. ASVs with greater than *>*0.01% abundance are plotted. Specific ASVs for BpSCSK (orange) and BpKH6 (blue) are plotted below. Error bars indicate standard deviation. **c,** Average number of unique ASVs detected. Error bars indicate 95% confidence intervals. **d,** Average Shannon diversity. Error bars indicate 95% confidence intervals. **e,** Bray-Curtis dissimilarity principal coordinates analysis (PCoA) for all time points.

The 16S rRNA gene sequencing revealed that, similar to the FMT, co-housing BpSCSK-colonized mice with SPF mice did not overcome the barrier to colonization established by BpSCSK. Over the two weeks, BpSCSK still remained approximately at 40% relative abundance **(Fig. 6b)**. Likewise, the number of unique ASVs and alpha diversity for BpSCSK-colonized mice remained at levels observed prior to co-housing **(Fig. 6c and 6d)**. The Bray-Curtis dissimilarity continued to show a clear separation from SPF mice as seen in previous experiments **(Fig. 6e)**. In addition to BpSCSK-treated mice, we sequenced fecal pellets from the SPF mice to determine whether co-housing would lead to colonization of BpSCSK. Although BpSCSK ASVs were detected in SPF mice, it was not consistently detected over the course of the two weeks **(Fig. 6b)**. Therefore, it could not be determined whether BpSCSK colonized or transiently passed through the gastrointestinal (GI) tract following coprophagia.

In conjunction with the continued lack of diversity, there were no notable changes in quantified metabolites between pre-co-housing and post-co-housing in BpSCSK-colonized mice except for alloisolithocholic and 3-oxolithocholic acid detected on the final day of collection at day 58 **(Extended Data Fig. 6a and 6b)**. Similarly, there were no major changes among quantified metabolites for SPF mice after two weeks. Thus, even an FMT followed by co-housing was unable to restore important microbiota-derived metabolites in the presence of BpSCSK.

### BpSCSK-resistant microbiota can restore gut diversity and key metabolite producers

BpSCSK was isolated from MyD88^-/-^ C57BL/6 mice that had been continuously treated with Augmentin, a combination of the *β*-lactam amoxicillin and *β*-lactamase inhibitor clavulanate, in drinking water for over 10 years^30^. This treatment has led to a significant shift in the microbiota composition compared to SPF mice. However, the MyD88^-/-^ C57BL/6 mice established a diverse microbiota, known as the ampicillin-resistant microbiota (ARM), with *α*-diversity comparable to SPF mice **(Fig. 3c and 3d; Extended Data Fig. 7c and 7d)**. ARM has fewer Bacillota and more species in the Bacteroidales order, especially with ASVs for *Muribaculaceae*, *Bacteroidaceae*, and *Porphyromonadaceae* **(Extended Data Fig. 7b)**. Since BpSCSK is in ARM, we hypothesized that the gut commensals from these mice would be capable of colonizing BpSCSK-dominated mice. To test this, we co-housed MyD88^-/-^ C57BL/6 mice harboring the ARM with BpSCSK-colonized mice and sequenced fecal pellets over the following two weeks **(Extended Data Fig. 7a)**. To reduce the impact of bacteria that were not associated with the ARM from colonizing, all mice were treated with ampicillin in drinking water throughout the experiment. Because BpSCSK is sensitive to ampicillin, we gavaged the previously described 4-mix consortia consisting of *Enterocloster* [*Clostridium*] *bolteae*, *Phocaeicola* [*Bacteroides*] *sartorii*, BpSCSK, and *Parabacteroides distasonis* (CBBP)^30,31^. *P. distasonis* and *P. sartorii* encode *β*-lactamases that degrade ampicillin.

As expected, the ARM containing BpSCSK was able to transfer and colonize mice treated with ampicillin and subsequently given the CBBP consortia **(Extended Data Fig. 7b)**. By day 14 of co-housing, the CBBP mice reached similar levels of unique ASVs as the ARM **(Extended Data Fig. 7c)**. Additionally, Shannon diversity was no longer significantly different between the two groups **(Extended Data Fig. 7d)**. Although BpSCSK was more abundant than in ARM, the average relative abundance of BpSCSK decreased from 41% at day 0 to approximately 18% by day 10 and 14 **(Extended Data Fig. 7b)**. Importantly, metabolomic profiling revealed that 5-aminovalerate and butyrate were detectable in CBBP-colonized mice by day 6 **(Extended Data Fig. 8a)**. Furthermore, proline levels decreased over the two-week period. While primary and secondary bile acids fluctuated between mice, some mice were able to modify bile acids **(Extended Data Fig. 8b)**. Thus, a microbiota developed in the presence of BpSCSK can restore diversity, including butyrate and 5-aminovalerate production, consumption of proline, and bile acid modification.

### Mice colonized with BpSCSK are susceptible to infection

Lantibiotics are active against Gram-positive bacteria and less effective against Gram-negative bacteria due to their outer membrane. Furthermore, butyrate depletion in the lower GI tract is associated with increased oxygen concentrations at the epithelial barrier, which can provide facultative anaerobes with a growth advantage^39^. Therefore, we tested the susceptibility of BpSCSK-colonized mice to intestinal colonization with *Klebsiella pneumoniae* MH258, a multi-drug resistant strain originally isolated from the blood of a patient undergoing cancer treatment^40^.

Mice pre-treated with ampicillin and gavaged with PBS or BpSCSK were challenged 34 days later by gavage with 1000 colony-forming units (CFUs) of *K. pneumoniae* MH258 **(Fig. 7a)**. For the positive control, SPF mice were treated for 4 days with ampicillin starting 6 days prior to challenge. Based on 16S rRNA gene sequencing and CFUs for *K. pneumoniae* MH258, BpSCSK-colonized mice were highly colonized with *K. pneumoniae* MH258 colonization than PBS-treated mice **(Fig. 7b)**. Importantly, although SPF + ampicillin-treated mice had greater relative abundance and CFUs of *K. pneumoniae* MH258 at day 1, BpSCSK-treated mice consistently maintained greater than 1 × 10^8^ CFU per gram of feces at day 1, 3, and 7 post infection **(Fig. 7c)**. In contrast, SPF + ampicillin-treated mice had reduced *K. pneumoniae* MH258 CFUs by day 3 that further decreased 7 days post infection.

**Fig. 7.**
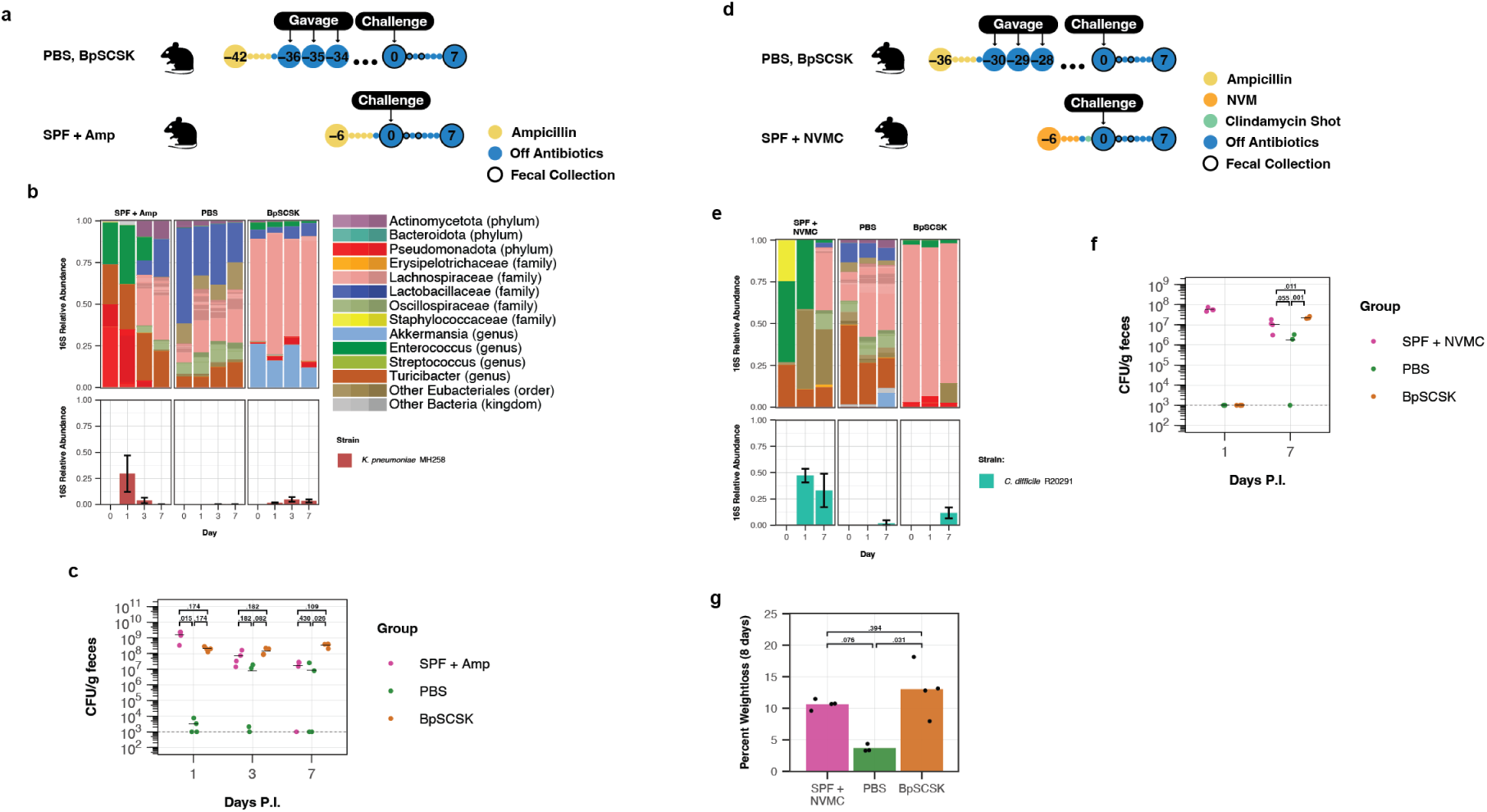
Mice colonized with BpSCSK are susceptible to infection. **a,** Schematic of experimental groups and time points. Female wild-type SPF C57BL/6 mice 9-10 weeks old were treated with 0.5 g/L ampicillin (Amp) in their drinking water for 4 days and given PBS or BpSCSK via oral gavage. On day 0, mice were challenged with approximately 1000 CFU of *K. pneumoniae* MH258 via oral gavage. Fecal samples were collected at indicated time points. Each group contains 4 mice. **b,** Average fecal microbiota 16S rRNA gene relative abundance. ASVs with greater than *>*0.01% abundance are plotted. Specific ASVs for *K. pneumoniae* MH258 are plotted below. Error bars indicate standard deviation. **c,** Colony-forming units (CFU) for *K. pneumoniae* MH258 post infection (P.I.). Black bars indicate the mean. Dashed line indicates the limit of detection. Statistical comparisons between individual groups were analyzed using Kruskal-Wallis test or Welch’s ANOVA followed by the Dunn’s or Welch t test for post-hoc comparisons. Numbers above the bars are the adjusted p-values. **d,** Schematic of experimental groups and time points. Female wild-type SPF C57BL/6 mice 9-10 weeks old were treated with 0.5 g/L Amp in their drinking water for 4 days and given PBS or BpSCSK via oral gavage. Positive control mice were treated with neomycin, vancomycin, and metronidazole in the drinking water followed by an intraperitoneal injection of clindamycin. On day 0, mice were challenged with approximately 200 *C. difficile* R20291 spores via oral gavage. Fecal samples were collected at indicated time points. Each group contains 3-4 mice. **e,** Average fecal microbiota 16S rRNA gene relative abundance. ASVs with greater than *>*0.01% abundance are plotted. Specific ASVs for *C. difficile* R20291 are plotted below. Error bars indicate standard deviation. Statistical comparisons between individual groups were analyzed using an ANOVA followed by the TukeyHSD post-hoc test. Numbers above the bars are the adjusted p-values. **f,** CFU for *C. difficile* R20291 P.I.. Black bars indicate the mean. Dashed line indicates the limit of detection. Numbers above the bars are the adjusted p-values. **g,** Average maximum percentage of weight loss that mice experienced over the 8 days mice were monitored starting at day 0 P.I. Each dot represents an individual mouse. Statistical comparisons between individual groups were analyzed using the Kruskal-Wallis test followed by the Dunn post-hoc test.

Bile acids are extensively modified by commensal bacteria into forms that can enhance or inhibit pathogen growth in the gut. Taurocholic acid, for example, induces germination in *Clostridioides difficile* while secondary bile acids can inhibit germination and vegetative growth^41^. *C. difficile* requires proline and glycine as reductive substrates to carry out Stickland fermentation for energy production and growth, and thus commensals can inhibit *C. difficile* by depleting these amino acids^42^. *In vitro* minimum inhibitory concentration (MIC) testing of blauticin by Zhang et al. demonstrated that *C. difficile* VPI10463 had a higher MIC compared to other gut commensals^17^. Thus, the metabolomic profile of BpSCSK-colonized mice and potential resistance to concentrations of blauticin in the gut indicate that BpSCSK may maintain a permissive environment for *C. difficile* infection weeks after antibiotic treatment.

Infection and disease susceptibility were tested by challenging mice with *C. difficile* R20291 28 days following oral gavage of PBS or BpSCSK. Fecal samples were collected and sequenced 0, 1, and 7 days post infection **(Fig. 7d)**. SPF mice given an antibiotic cocktail of neomycin, metronidazole, vancomycin, and clindamycin (NMVC) prior to the challenge served as a positive control for infection. Although 16S rRNA gene sequences for *C. difficile* could be detected in some mice from all groups by day 7 post infection, all mice in the SPF + NMVC and BpSCSK-treated groups were colonized by *C. difficile* **(Fig. 7e)**. This was further supported by quantifying the CFUs of *C. difficile* R20291 in the fecal pellets **(Fig. 7f)**. Interestingly, over the 8 days that mouse weight was monitored post infection, BpSCSK-colonized mice showed similar weight loss to the positive control mice relative to the starting weight when challenged **(Fig. 7g)**. On average, PBS-treated mice had less than a 5% loss from their initial weight. Consistent with proline and glycine reductase mediated Stickland fermentation in *C. difficile* R20291, SPF + NVMC and BpSCSK-treated mice exhibited decreased proline and glycine upon *C. difficile* R20291 infection **(Extended Data Fig. 9a)**. Additionally, the concentration of 5-aminovalerate, the by-product of Stickland fermentation with proline, was increased.

These findings indicate that BpSCSK-colonized mice are vulnerable to colonization by pathogens like *K. pneumoniae* and *C. difficile*, remaining susceptible for months after cessation of antibiotic treatment.

## Discussion

Bacteria have evolved strategies to compete with one another, such as nutrient competition, predation, and bacteriocin production. While narrow- and broad-spectrum bacteriocins have been identified, much is still unknown about their overall impact on microbiome compositions and functions. Lantibiotics and lantibiotic-producing bacteria are being used in the food industry, and they are being investigated for their potential to combat pathogens that have acquired resistance to clinically available antibiotics. Previous studies have demonstrated that purified lantibiotics can impact gut commensals^18,22,23^, and lantibiotic-producing bacteria have been shown to inhibit human gut commensals *in vitro* ^19^. Since bacteriocin-producing bacteria are being investigated as an approach to combat increased antibiotic resistant pathogens, it is becoming increasingly important to understand their influences on the microbiome.

Analyses of fecal samples collected from healthy individuals and hospitalized patients uncovered a high prevalence of lanthipeptide-encoding genes in their intestinal microbiomes, including previously characterized lantibiotics. While activity, specificity, and expression of lanthipeptides vary widely, we found certain lanthipeptides correlate with a fitness advantage for the encoding species, leading to greater relative abundance in the microbiota. The presence of identical lanthipeptide genes across different patients may indicate that patients are coming into contact with common bacterial sources. Some genera, such as *Blautia*, can form spores, and be acquired from the environment. Early colonization with a lantibiotic or other potent bacteriocin-producer, in a developing or dysbiotic microbiota, has the potential to impair microbiota diversification. Importantly, many of the species capable of producing lantibiotics overlap with species that colonize the infant gut microbiome in early life^43^. In these and other settings, bacteria expressing lantibiotics may serve as “pathogen facilitators”, i.e. microbes that do not directly cause disease but establish a state of dysbiosis that enables pathogen persistence. Identifying narrower spectrum lantibiotics that resist pathogens while maintaining beneficial commensal bacterial species represents a significant challenge.

In this study, we demonstrate that BpSCSK, a strain of *Blautia* with a known lantibiotic-producing BGC, alters recolonization of the gut microbiota in mice with antibiotic-induced dysbiosis. Our previous studies demonstrated that BpSCSK culture supernatants inhibited Grampositive but not Gram-negative commensal bacterial strains^31^. However, using an *in vivo* mouse model, our results demonstrate that many Gram-positives and Gram-negative commensal bacteria are suppressed by BpSCSK gut colonization. This may result from inhibition of critical species that provide the nutrients or other factors that enable colonization by broader commensal populations. Additionally, BpSCSK prevents recolonization by butyrate producers, which can lead to inhibition of oxygen-sensitive commensal strains in the colonic lumen^39^. Together, these provide possible explanations for the inhibition of bacteria that may otherwise be resistant to blauticin lantibiotics. Given that BpKH6 did not completely restrict recolonization, direct nutrient depletion by *Blautia* likely presents only a partial mechanism of commensal suppression. Since BpSCSK was unable to colonize SPF mice that were not pre-treated with ampicillin, it is likely that early colonization and expansion is required to resist colonization by other commensals.

Hindering gut recolonization with key-metabolite producers leads to prolonged suppression of metabolites that support a healthy gut environment. Despite BpSCSK’s ability to provide colonization resistance against VRE, our study reveals that it enhances colonization and infection with *K. pneumoniae* and *C. difficile*. This finding highlights the importance of characterizing commensal bacterial strains that might be included in live biotherapeutic products designed to reestablish microbiome functions in patients with dysbiosis. Avoiding undesirable side effects such as prolonged dysbiosis and susceptibility to additional pathogens is critical. In addition to increasing susceptibility to pathobiont infections, BpSCSK also reduced FMT-mediated reconstitution with commensal microbes and thus might reduce its effectiveness at treating recurrent *C. difficile*. As a result, key microbiota-derived metabolites remained below the limits of detection despite the attempts to restore the microbiota.

Suez et al., using an *in vivo* murine model and human volunteers, have shown that bacterial strains used as probiotics are capable of prolonging antibiotic-induced dysbiosis^29^. While a mechanism was not established, they found that the supernatant from the probiotics inhibits bacteria cultured from human feces. The same species used in their probiotic treatment were identified in our study to contain lanthipeptide genes. In addition, they found that *Blautia* could grow in the presence of the probiotic strains, which could be the result of lantibiotic resistance genes that often accompany lantibiotic-producing bacteria. Although more work needs to be done to establish a connection, our findings suggest a potential mechanism for the prolonged dysbiosis caused by some probiotics.

Despite the negative impact of BpSCSK on microbiota resilience following antibiotic treatment, lantibiotic-producing strains may still find a role in reducing infections caused by highly antibiotic-resistant pathogens. Antibiotic resistant pathogens, such as VRE, colonize the gut at high density and can lead to systemic infections that are becoming increasingly resistant to the few remaining antibiotic options. Intestinal clearance of VRE with BpSCSK followed by antibiotic-mediated clearance of BpSCSK represents a potential two-step path to reducing the risk of system infection with nearly untreatable pathogens. More work is needed to determine the therapeutic potential of lantibiotics and to better characterize the novel lantibiotics currently found in probiotics and microbiomes to mitigate off-target side-effects that could impair commensal bacterial functions.

## Experimental procedures

### Human fecal sample collection

Fecal samples for clinical patients were collected by nurses in inpatient wards and medical intensive care units. Upon collection, samples were immediately sent for storage at 4 °C via a pneumatic tubing system. The samples in storage at 4 °C were then aliquoted and stored at −80 °C within 24 hours after initial collection.

### Shallow shotgun sequencing

Human fecal samples were suspended in lysis buffer and broken up with a bead beater (BioSpec Product). DNA was purified with the QIAamp PowerFecal Pro DNA kit (Qiagen). Sequencing was performed using the Illumina HiSEq platform. This produced approximately 7-8 million pairedend reads for each sample with a read length of 150 bp. After trimming adapters from raw reads, the quality of reads was assessed and controlled with Trimmomatic^44^ (version 0.39). Reads that map to the human genome were removed with kneaddata (version 0.7.10). In order to assign taxonomy, the filtered reads were run through MetaPhlAn4^36^. All reads were then assembled into contigs using MEGAHIT^45^ (version 1.2.9).

### 16S rRNA gene sequencing

Mouse fecal pellets were collected in 2 mL cryogenic microcentrifuge tubes (Sarstedt) on collection days and placed on dry ice until transferred to a −80 °C freezer until sequenced. Fecal samples were suspended in lysis buffer and broken up with a bead beater (BioSpec Product). DNA was purified with the QIAamp PowerFecal PRo DNA kit (Qiagen). With the purified DNA, the V4-V5 region of the 16S rRNA gene was amplified using universal primers 563F (5‘-nnnnnnnn-NNNNNNNNNNNN-AYTGGGYDTAAA-GNG-3’) and 926R (5‘-nnnnnnnn-NNNNNNNNNNNN-CCGTCAATTYHT-TTRAGT-3’), where ‘N’ represents the barcodes and ‘n’ are additional nucleotides added to offset primer sequencing. Amplicons containing approximately 412 bp regions were purified in a spin column (Minelute, Qiagen). Purified amplicons were quantified using a Qubit 2.0 fluorometer and pooled at equimolar concentrations. Unique Dual Index adapters were ligated to the amplicons with QIAseq 1-step amplicon library kit (Qiagen). Library quality control was performed using Qubit and TapeStation. Results were analyzed as using the Phyloseq^46^ R package. Taxonomy for amplicon sequence variants (ASVs) were assigned with Ribosomal Database Project (RDP) classifier 2.14^47^. *α*-diversity was calculated using the estimate richness() function, Bray-Curtis dissimilarity was calculated and plotted in R using the ordinate() and plot ordination() functions.

### 16S rRNA gene qPCR

An aliquot of the samples prepared for 16S rRNA gene sequencing were diluted to 20 ng/µL. The primers 563F (5’-AYTGGGYDTAAAGNG-3’) and 926R (5’-CCGTCAATTYHTTTRAGT-3’) for the V4-V5 region of the 16S rRNA gene were used for amplification. A standard curve was created with a V4-V5 region on a linearized TOPO pcr2.1TA vector isolated from *Escherichia coli* DH5*α* cells. The vector was serially diluted 5-fold ranging from 10^8^ to 10^3^ copies/µL. qPCR was performed on a QuantStudio 6 Pro (Applied Biosystems) using PowerTrack SYBR Green Master Mix (A46109) with the following cycling conditions: 95 °C for 10 min, followed by 40 cycles of 95 °C for 30 s, 52 °C for 30 s,and 72 °C for 1 min. Design and Analysis v2 software was used to calculate the copy numbers for each sample. Copy numbers were normalized based on the weight of fecal pellets the DNA was extracted from.

### Lanthipeptide detection

Anvi’o^48^ was used to manage gene annotations. Anvi’o contig databases were generated from fasta files containing shotgun sequencing reads assembled into contigs by MEGAHIT^45^ (version 1.2.9) using ‘anvio-gen-contigs-database.’ Genes in the Anvi’o contigs databases were called using Prodigal^49^ (version 2.6.3) with default parameters. Specific Pfam domains^35^ were annotated using ‘anvi-run-hmms’ with custom Hidden Markov Model (HMM) profile^50^. Contigs were also run through the RODEO (Rapid ORF Description and Evaluation Online)^51^ program to identify lanthipeptide genes. All contigs that contained an annotated Pfam domain, previously identified lanthipeptide from Walker et al.^7^, a lanthipeptide gene identified by RODEO, or lanthipeptides in the InterPro^52^ and KEGG^53^ databases were extracted and converted into a new fasta file. The contigs were then made into a new Anvi’o contig database and annotated with ‘anvi-run-pfams.’ Additionally, the fasta file containing filtered contigs was run separately through Bakta^54^ and genes were extracted from the Anvi’o contig database to run through GhostKOALA^55^ for gene annotations. Any newly identified lanthipeptides from the gene annotations detected were added to the list of lanthipeptides and were re-screened to identify any additional contigs containing the gene. Genes of interest were also run through NCBI BLASTp to further identify and confirm lanthipeptide sequences^56^. Only the top 5 hits from BLASTp were considered. Lanthipeptide class was assigned based on annotations or surrounding synthesis genes.

### Taxonomic classification

Taxonomic classification for contigs containing lanthipeptides was determined by running contig sequences through the NCBI Nucleotide database^56^. Hits were limited to the top five results. A single taxonomic classification was assigned to a contig if the evalue was less than 1 × 10^−3^, percent identity was greater than 75%, and query coverage was greater than 60%. If there were more than one result remaining, the result with the highest bitscore was used. Contigs that did not have a taxonomic classification that met the thresholds were labeled “Unclassified.”

### Lanthipeptide fitness advantage assessment

Taxonomic annotations for contigs were matched to identical genus and species taxonomic classifications determined by MetaPhlAn4^36^. For species that were annotated as “sp.”, the taxids were appended to the taxonomic classification for matching purposes. Only lanthipeptides that had at least five matches were considered. Relative abundances of species that did not contain a match to a lanthipeptide were averaged together. Fold changes were calculated as the ratio of the relative abundances of bacteria that contained the lanthipeptide genes to the average relative abundances of the same species that did not contain the lanthipeptide gene. Next, the fold changes of lanthipeptides were averaged together to get the mean fold change for each lanthipeptide gene. Results were plotted as the log_2_ mean fold change.

### UMAP analysis of lanthipeptides

Lanthipeptide sequences were aligned in R using MUSCLE^57^ in the msa^58^ R package. A distance matrix as generated using dist.alignment() in the seqinr^59^ package. Next, the uniform manifold approximation and projection (UMAP)^60^ conducted using the umap R package.

### Mice

All mouse studies were approved by The University of Chicago Institutional Animal Care and Use Committee (protocol 72599). Wild-type C57BL/6 mice were ordered from the AX8 breeding room at Jackson Laboratories. MyD88^-/-^ C57BL/6 mice are maintained as a colony at the University of Chicago under continuous treatment of augmentin (NorthStarx NDC:16714-293-01), which contains 0.48 g/L amoxicillin and 0.07 mg/L clavulanate, added to drinking water. For experiments, female mice aged 9 to 10 weeks old and 37 to 44 weeks old for wild-type and MyD88^-/-^ mice, respectively. All mice were housed in a BSL2 animal room under specific-pathogen-free (SPF) conditions at the University of Chicago. Mice were fed a standard chow diet and given autoclaved acidified drinking water. All mice were randomized and single housed in cages, except where specified, containing corncob bedding. Cages were changed every 2 weeks. When working with the mice, a 10% bleach solution was used to disinfect surfaces to prevent contamination from spores.

### Bacterial consortia strains and growth conditions

Frozen stocks for *Enterocloster bolteae* CBBP, *Phocaeicola sartorii* CBBP, *Parabacteroides distasonis* CBBP, *Blautia pseudococcoides* SCSK, and *Blautia producta* KH6 were streaked out on Columbia blood agar plates with 5% sheep’s blood (BBL 221263) and incubated at 37 °C in an anaerobic chamber for 2 days. Single colonies were mixed in 500 µL of Dulbecco’s phosphate buffered saline (DPBS) (Gibco 14190-144) and 200 µL were streaked in a lawn on a fresh plate. Plates were again incubated for 2 days. Next, lawns were scrapped up with a sterile inoculating loop and suspended in 1.2 mL DPBS. For the CBBP consortia, each strain was mixed into the same 1.2 mL DPBS. 200 µL was administered via oral gavage for colonization in mice. These steps were repeated for each day the bacteria were administered.

### FMT and Co-housing

For the FMT, one fecal pellet was collected from four female SPF C57BL/6 mice. The fecal pellets were pooled into one tube and kept on ice until transferred to an anaerobic chamber within 30 minutes from collection. All four fecal pellets were combined and suspended in 4 mL of reduced DPBS. Then, the fecal pellets were crushed, mixed, and aliquoted in 200 µL volumes for gavage. Mice were gavaged with 200 µL. This process was then repeated on subsequent days for a total of three gavages. Mice used for the FMT control were treated with ampicillin (Athenex NDC:70860-118-99) for four days starting six days before the first gavage. After two weeks, each mouse colonized with BpSCSK was then co-housed with a female SPF C57BL/6 mouse for an additional two weeks. Fecal pellets were collected for metagenomic and metabolomic analysis throughout the experiments.

In a separate experiment, BpSCSK colonized mice were individually co-housed with female MyD88^-/-^ C57BL/6 mice. All mice were kept on 0.5 g/L ampicillin in the drinking water throughout the experiment. Fecal pellets were collected for metagenomic and metabolomic analysis.

### Antibiotic treatment

Mice treated with ampicillin to deplete the native gut microbiota were given 0.5 g/L ampicillin in their drinking water starting 6 days prior to their first gavage and returned to their normal drinking water two days before their first gavage. MyD88^-/-^ and mice colonized with the CBBP consortia were continuously treated with 0.5 g/L ampicillin prepared every 4 days. Mice treated with a cocktail of antibiotics were given 0.25 g/L neomycin (Fisher BioReagents BP2669), vancomycin (SAGENT Pharmaceuticals NDC:25021-158-99), and metronidizole (Sigma-Aldrich M3761) in their drinking water starting five days from infection with *C. difficile*. They were returned to normal drinking water 2 days prior to infection and given an intraperitoneal (IP) injection of 100 µL of 2 g/L clindamycin (Sigma-Aldrich C5269) 24 hours prior to infection.

### *C. difficile* spore preparation

*C. difficile* spore preparation was carried out as previously described with slight modifications^61^. Using isolated colonies grown on BHIS agar, which consists of 37 g/L Brain-Heart infusion powder (BD Bacto 237500), 5 g/L yeast extract (BD Bacto 212750), 15 g/L agar (BD Bacto 214040), and 0.1% (w/v) L-cysteine powder (Sigma cat# C7352), *C. difficile* was inoculated into prereduced BHIS broth and incubated anaerobically in a 50 mL conical tube at 37 °C. After 40-50 days, the suspended cells were harvested via centrifugation with five washes with ice-cold water. The cell pellet was then re-suspended in 20% HistoDenz (w/v) (Sigma, St. Louis, MO) and layered onto a 50% (w/v) HistoDenz solution. Purified spores were collected via centrifugation at 15,000 g for 15 min 3 times. The resulting spore pellet was again washed with ice-cold water 4 times to remove traces of HistoDenz. Finally, the spore pellet was re-suspended in sterile water. To kill any remaining vegetative cells, the re-suspended spores were heated on a heat block for 20 minutes at 60 °C. A portion of the spore stocks were diluted and plated on BHIS agar to confirm that there is less than 1 vegetative cell per 200 spores. Another portion was plated on BHIS agar containing 0.1% (w/v) taurocholic acid (BHIS-TA) for numeration.

### C. difficile infection

Mice were challenged via oral gavage of approximately 200 *C. difficile* R20291 spores suspended in 200 µL of DPBS. Fecal samples were collected prior to challenge followed by day 1 and 7 post infection for CFU counts, 16S rRNA gene sequencing, and metabolomics. Mice were also weighed daily for 8 days and compared to their weight prior to challenge.

### K. pneumoniae infection

*K. pneumoniae* MH258 was grown aerobically at 37 °C overnight in LB broth supplemented with 100 µg/mL carbenicillin (Fisher BioReagents BP2648) and 50 µg/mL neomycin. Overnight culture was serial diluted to 5 CFU/µL in DPBS. Mice were challenged via oral gavage of approximately 1000 CFU (200 µL) of *K. pneumoniae*. Fecal samples were collected prior to infection in addition to day 1, 3, and 7 post infection for CFU and 16S rRNA gene sequencing.

### Quantification of *C. difficile* and *K. pneumoniae* shedding

Fecal pellets were collected from mice on the indicated dates in micro-centrifuge tubes and stored on ice. After weighing samples, 1 mL of DPBS was added to each tube and fecal pellets were broken up using an inoculating loop. Then, samples were vortexed for 5 seconds and aliquoted to a 96-well flat-bottom plate for serial dilutions. Each sample was serial diluted in DPBS to 10^−5^ in 10-fold steps using a total of 300 µL volumes for each dilution. Next, 10 µL of each dilution was plated in rows on agar media and incubated at overnight at 37 °C within 1 hour of collection. For *C. difficile*, serial dilutions and culturing were conducted in an anaerobic chamber and plated on CC-BHIS-TA containing 0.25 g/L D-cycloserine (Acros Organics 228480250), and 0.016 g/L cefoxitin (Sigma C4786). *K. pneumoniae* was grown on LB (BD 244620) agar containing 100 g/mL carbenicillin, 50 g/mL neomycin, and 25 g/mL chloramphenicol (Fisher BioReagents BP904) antibiotics. Colonies were counted at the least diluted concentration that colonies could be accurately counted at and reported as CFU per gram of feces.

### Quantitative metabolomics

Mouse fecal pellets were collected in 2 mL cryogenic microcentrifuge tubes (Sarstedt 72.694.006) on collection days and placed on dry ice until transferred to a −80 °C freezer DNA sequencing. Fecal samples were weighed and extraction solvent, 80% methanol spiked with internal standards, was added to make a ratio of 100 mg of material per mL of extraction solvent. Samples were then homogenized at 4 °C using a Bead Mill 24 Homogenizer (Fisher; 15-340-163) set at 1.6 m/s with six x 30 s cycles, 5 seconds off per cycle. Next, samples were centrifuged at −10 °C, 20,000 x g for 15 min. The supernatant was used for subsequent metabolomic analysis.

Short-chain fatty acids (SCFAs), amino acids, and phenolic metabolites were quantified via derivatization with pentafluorobenzl-bromide (PFBBr). Methanol containing internal standards was added to fecal samples at 4 volumes per mg of feces. Samples were subjected to centrifugation followed by addition of 100 µL of 100 mM borate buffer at a pH of 10. Next, 400 µL 100 mM PFBBr suspended in acetonitrile was added to the samples and incubated at 65 °C for 1 hour. Lastly, metabolites were extracted with hexanes and subjected to gas chromatography-mass spectrometry (GC-MS) (Agilent 8890/5977B and 7890B/5977B) with chemical ionization and negative mode detection.

Bile acids were quantified via liquid chromatography-mass spectrometry (LC-MS). First, 75 µL was added to mass spectrometry autosampler vials (Microliter; 09-1200) and subsequently dried using a nitrogen stream of 30 L/min (top) and 1 L/min (bottom) at 30 °C (Biotage SPE Dry 96 Dual; 3579M). Dried samples were then re-suspended in 750 µL of a 50:50 water:methanol mixture. The vials were added to a thermomixer (Eppendorf) to re-suspend analytes under the following conditions: 4 °C, 1000 rpm for 15 min with infinite hold at 4 °C. To remove insoluble debris, samples were centrifuged in microcentrifuge tubes at 20,000 x g for 15 mins at 4 °C. Next, 700 µL was transferred to a new mass spectrometry autosampler vial and analyzed in negative mode on an LC system (Agilent 1290 infinity II) coupled to a quadrupole time-of-flight (QTOF) mass spectrometer (Agilent 6546) equipped with an Agilent Jet Stream Electrospray Ionization source. Then, 5 µL of sample was then injected onto an XBridge BEH C18 column (3.5 µm, 2.1 × 100 mm; Waters Corporation, PN) fitted with an XBridge BEH C18 guard (Waters Corportation, PN) at 45 °C. Elution started with 72% A (Water, 0.1% formic acid) and 28% B (Acetone, 0.1% formic acid) with a flow rate of 0.4 mL/min for 1 min and linearly increased to 33% B over 5 min, then linearly increased to 65% B over 14 min. Then the flow rate was increased to 0.6 mL/min and B was increased to 98% over 0.5 min and these conditions were held constant for 3.5 min. Finally, re-equilibration at a flow rate of 0.4 mL/min of 28% B was performed for 3 min. The electrospray ionization conditions were set with the capillary voltage at 3.5 kV, nozzle voltage at 2 kV, and detection window set to 100-1700 m/z with continuous infusion of a reference mass (Agilent ESI TOF Biopolymer Analysis Reference Mix) for mass calibration. A 10-point calibration curve was used for quantification. Data analysis was performed using MassHunter Profinder Analysis software (version B.10, Agilent Technologies) and confirmed by comparison with authentic standards. Normalized peak areas were calculated by dividing raw peak areas of targeted analytes by averaged raw peak areas of internal standards.

The log_2_ fold change for metabolites was calculated as the ratio of the average metabolite concentration for all mice within a group at a timepoint by the average SPF mouse concentrations across all timepoints.

### Statistical analysis

Statistical analyses were performed in R using the stats and rstatix packages. Normality was assessed using the Shapiro–Wilk test. Homogeneity of variance was evaluated with either Levene’s test (for non-normal data) or Bartlett’s test (for normal data). Depending on data distribution and variance assumptions, one of the following omnibus tests was used: ANOVA (normal data with equal variances), Welch’s ANOVA (normal data with unequal variances), or Kruskal–Wallis test (non-normal data). When the omnibus test was significant, appropriate post hoc tests were performed: Tukey’s HSD following ANOVA, Games–Howell or Welch’s t-test following Welch’s ANOVA, and Dunn’s test following Kruskal–Wallis. P-values were adjusted for multiple comparisons using the Benjamini–Hochberg method where appropriate.

## Acknowledgments

We thank the current and previous members of the Pamer lab for all their invaluable input on this manuscript. We thank the other members of Duchossois Family Institute especially the research staff that run the metagenomic and metabolomic core facilities. We also thank the Animal Resource Center at the University of Chicago for their maintenance and upkeep of the animal facilities used in this research. S.S.S was supported by the MSTP training grant T32GM150375.

## Author contributions

C.G.C. and E.G.P. designed the experiments, wrote, and edited the manuscript. S.R.D and D.A.M. performed RODEO analysis. N.P.D., H.L., C.K.W. and A.S. curated shotgun and 16S rRNA gene sequencing data and aided with lanthipeptide screening in clinical samples. N.P.D., H.L., and A.M.S. curated metabolomics data. E.J.M. maintained mouse colonies. C.G.C., Z.J.Z., Q.D., R.L.P., and S.S.S. performed experiments. C.G.C. performed data analysis. E.G.P. acquired funding.

---

**Extended Data Fig. 1.**
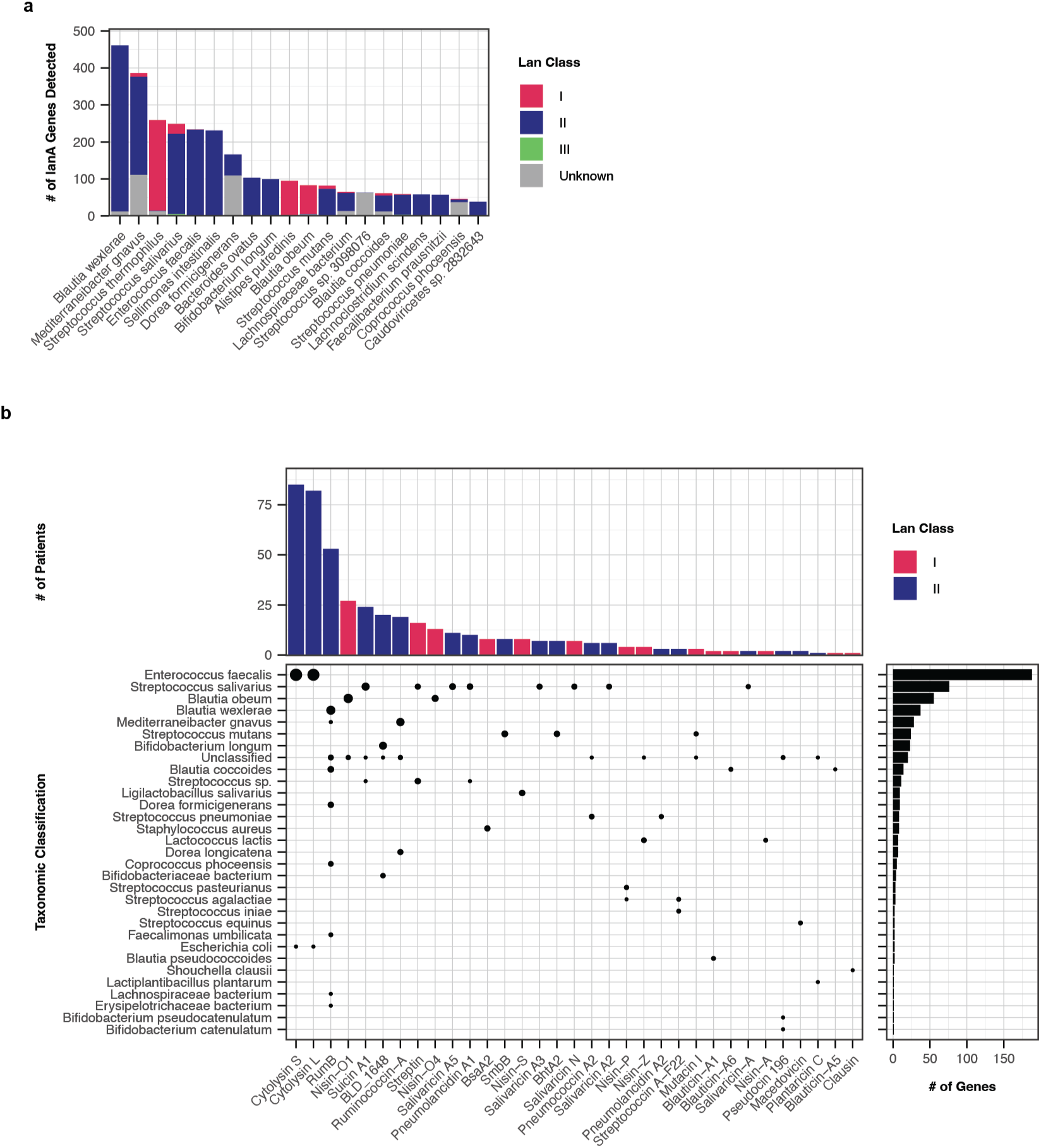
Species classification of lantibiotic-producers and previously characterized lanthipeptides in clinical patients. **a,** Distribution of detected lanthipeptide genes across the top 20 species assigned to the encoding contig. **b,** Lanthipeptides that matched the core sequence of previously characterized lanthipeptide sequences. Vertical columns represent the number of times the lanthipeptide was detected. Horizontal bars represent the number of lanthipeptides were found in a given species. The dots represent the intersection between the two plots with the size of the dots representing the number of times a specific lanthipeptide was found in a specific species.

**Extended Data Fig. 2.**
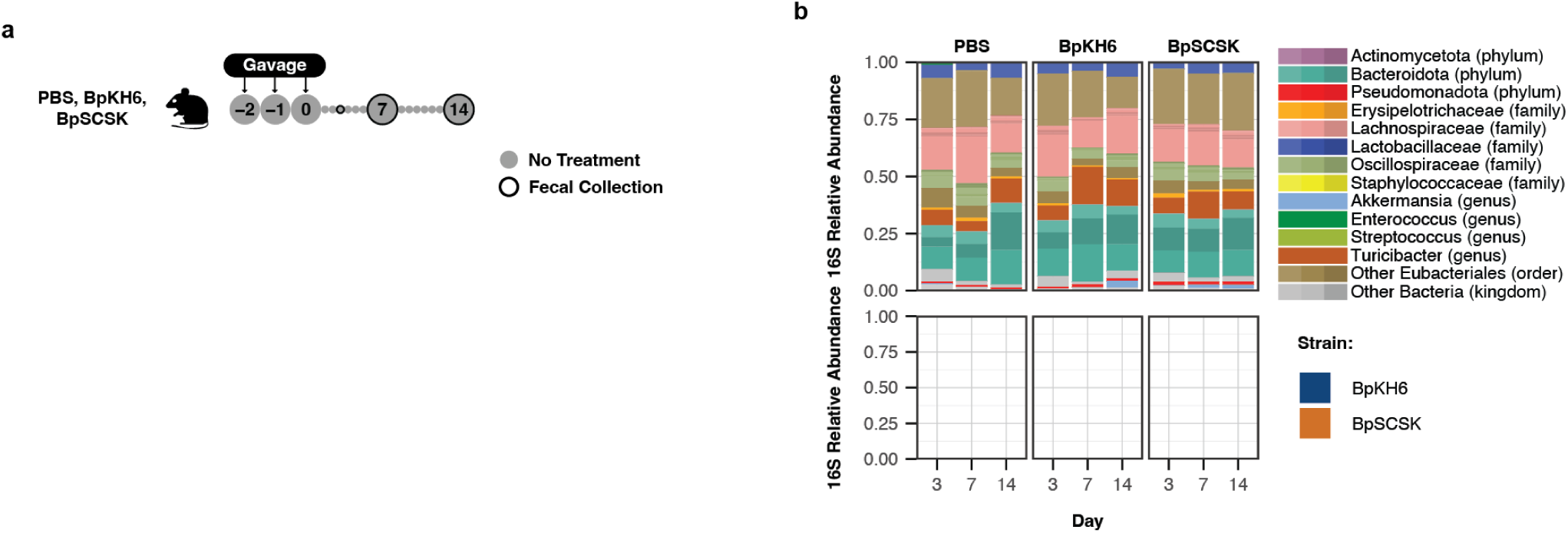
BpSCSK colonization in wild-type SPF mice. **a,** Schematic of experimental groups and time points. Female wild-type SPF C57BL/6 mice 9-10 weeks old were given PBS, BpKH6, or BpSCSK via oral gavage. Fecal samples were collected at indicated timepoints. Each group contains 3 mice. **b,** Average fecal microbiota 16S rRNA gene relative abundance. ASVs with greater than *>*0.01% abundance are plotted. Specific ASVs for BpSCSK and BpKH6 are plotted below.

**Extended Data Fig. 3.**
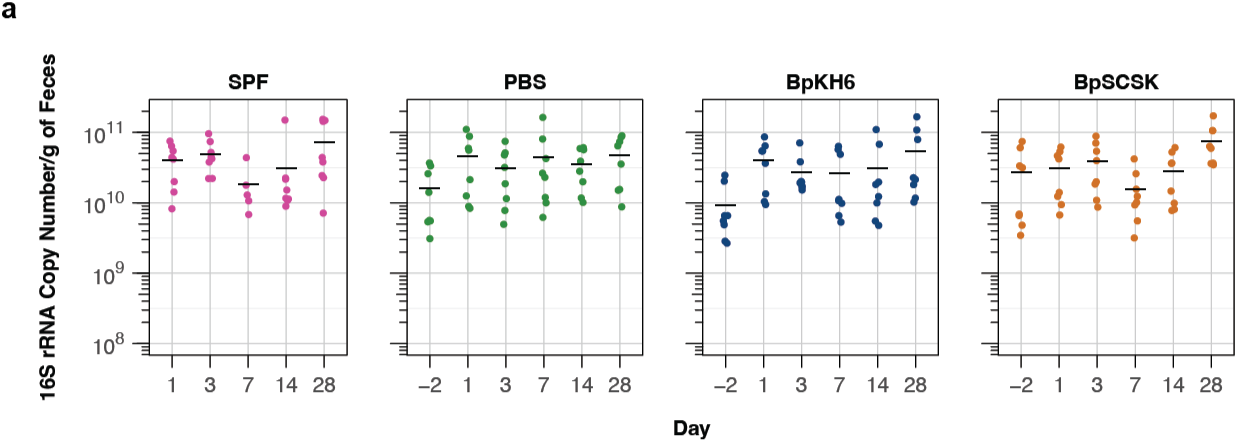
BpSCSK-treated mice have similar bacterial titers as SPF mice. **a,** 16S rRNA gene copy number per g of feces determined via qPCR. Black bars indicate the mean.

**Extended Data Fig. 4.**
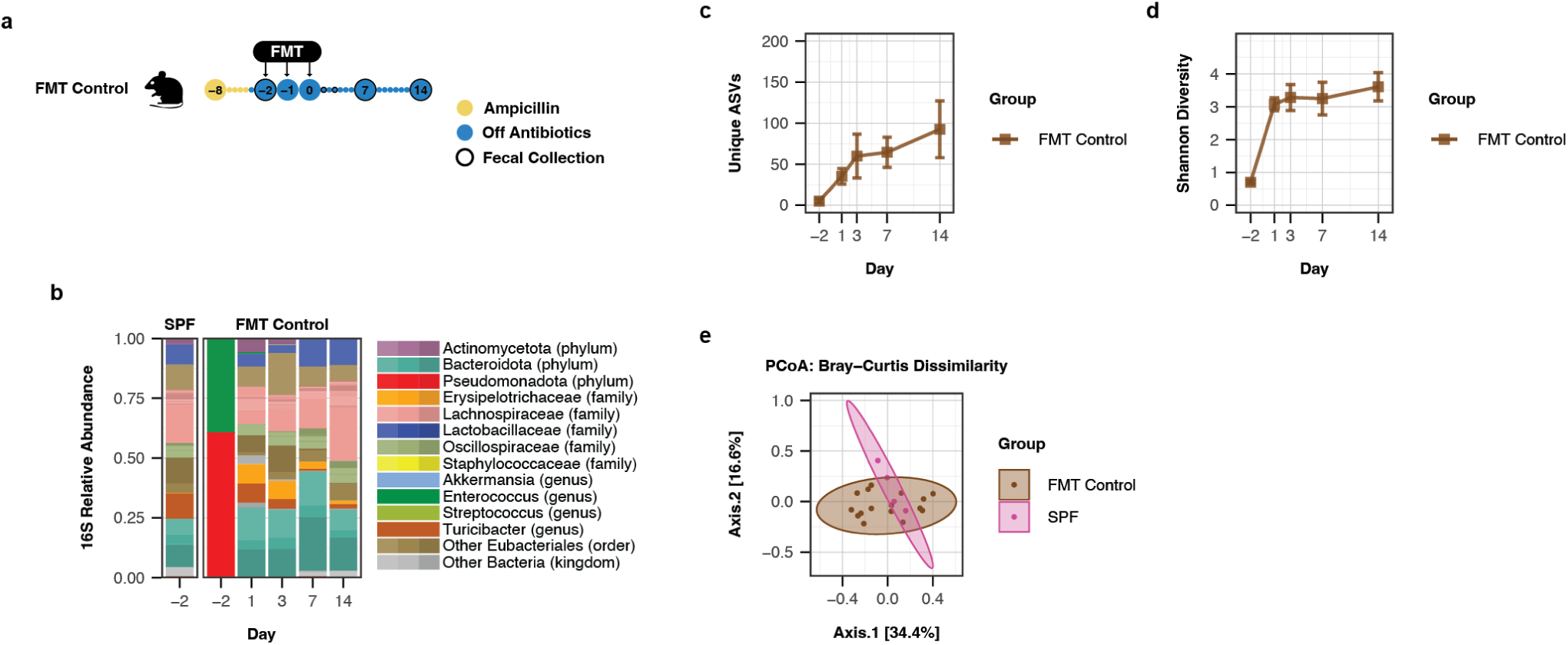
Colonization of FMT in antibiotic treated mice. **a,** Schematic of experimental groups and time points. 4 mice in the FMT control were subjected to an FMT from an SPF mouse for three days via oral gavage. The FMT Control group were subject to 4 days ampicillin treatment starting 6 days before the first FMT. **b,** Average fecal microbiota 16S rRNA gene relative abundance. ASVs with greater than *>*0.01% abundance are plotted. **c,** Average number of unique ASVs detected. Error bars indicate 95% confidence intervals. **d,** Average Shannon diversity. Error bars indicate 95% confidence intervals. **e,** Bray-Curtis dissimilarity principal coordinates analysis (PCoA) for all time points.

**Extended Data Fig. 5.**
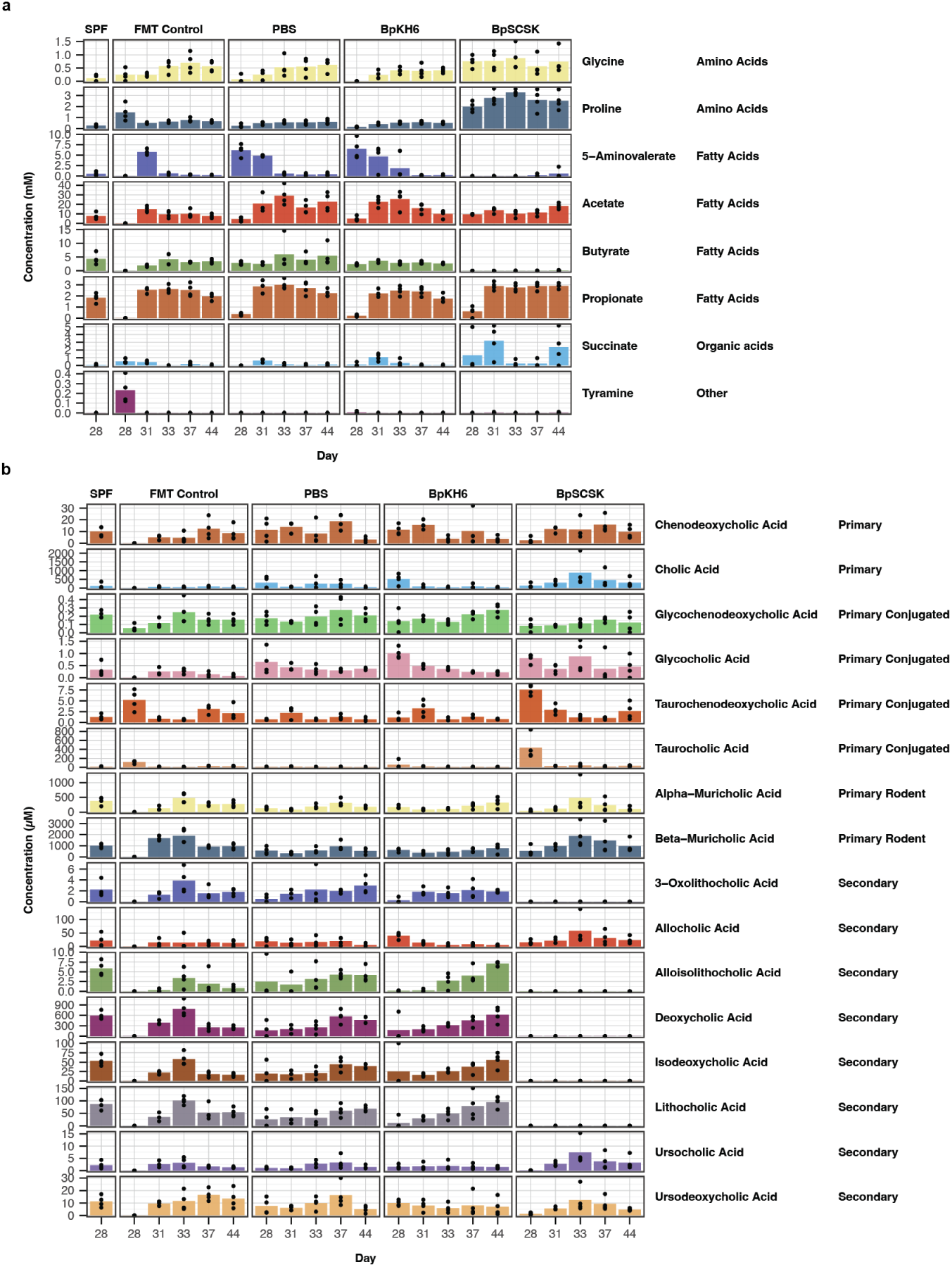
Metabolite concentrations after FMT treatment. a, b, Absolute concentrations of metabolites for mice from Fig. 5. The bars represent the average concentration across the mice on each day. The dots are the individual values for each mouse.

**Extended Data Fig. 6.**
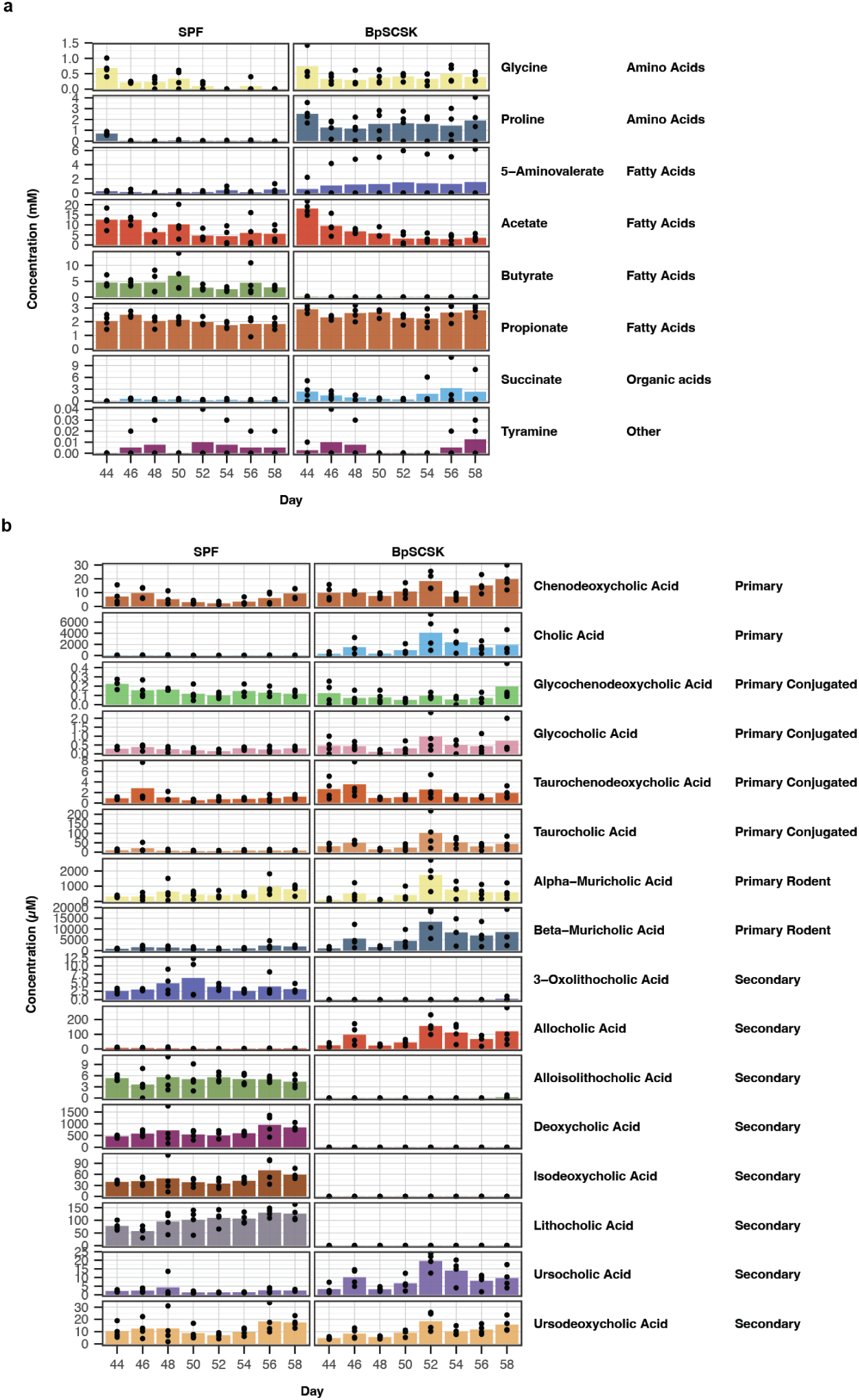
Metabolite concentrations after co-housing BpSCSK-colonized mice with SPF mice. a, b, Absolute concentrations of metabolites for mice from Fig. 6. The bars represent the average concentration across the mice on each day. The dots are the individual values for each mouse.

**Extended Data Fig. 7.**
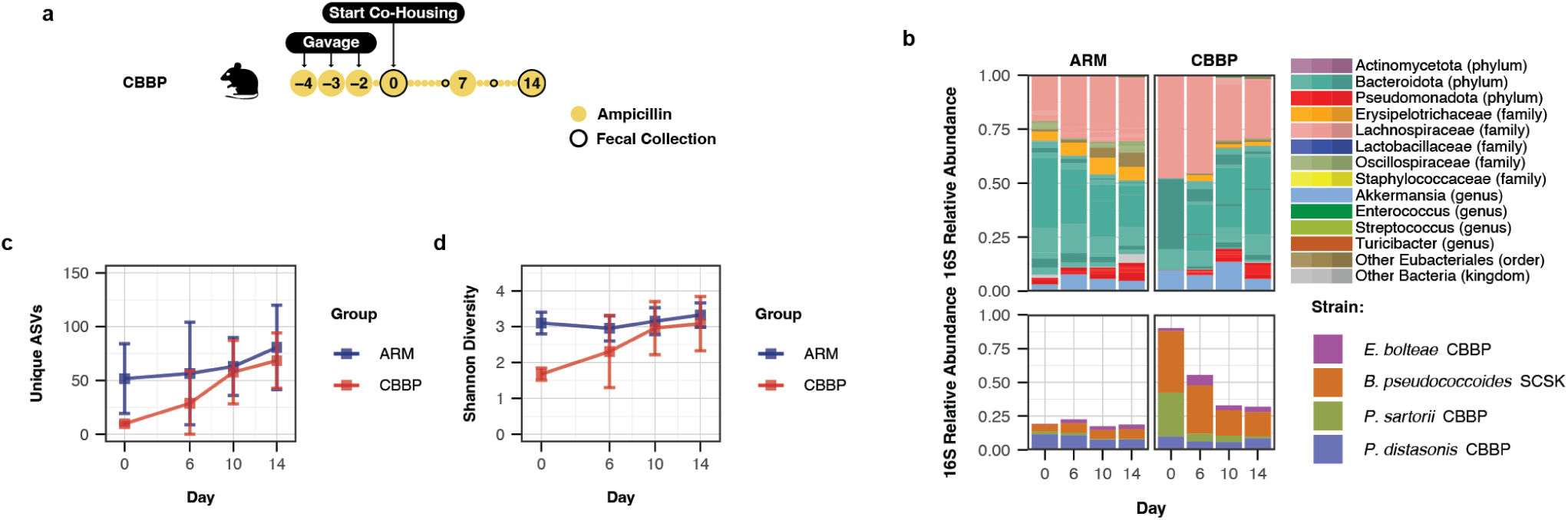
BpSCSK-resistant microbiota can restore gut diversity and key metabolite producers. **a,** Schematic of experimental groups and time points. 4 female C57BL/6 mice given CBBP (*Eubacteria bolteae* CBBP, *Phocaeicola sartorii* CBBP, *Blautia pseudococcoides* SCSK, and *Parabacteroides distasonis* CBBP) via oral gavage were co-housed with C57BL/6 MyD88-/- mice with an ampicillin resistant microbiota (ARM) for 2 weeks. Drinking water was supplemented with 0.5 g/L ampicillin throughout the experiment. **b,** Average fecal microbiota 16S rRNA gene relative abundance. ASVs with greater than *>*0.01% abundance are plotted. Specific ASVs for found in the CBBP consortium are plotted below. **c,** Average number of unique ASVs detected. Error bars indicate 95% confidence intervals. **d,** Average Shannon diversity. Error bars indicate 95% confidence intervals.

**Extended Data Fig. 8.**
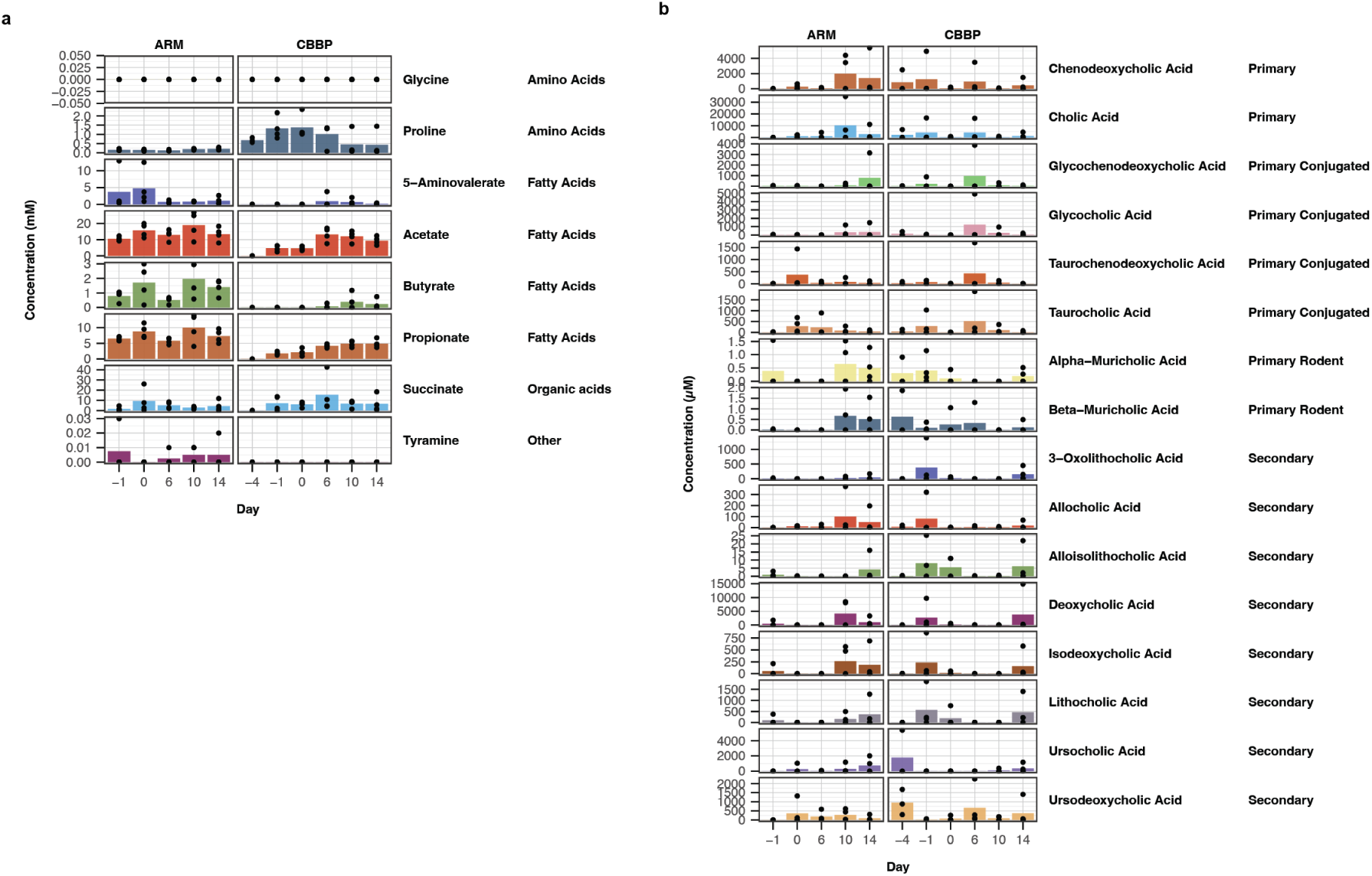
Metabolite concentrations after co-housing BpSCSK-colonized mice with ARM mice. a, b, Absolute concentrations of metabolites for mice from Extended Data Fig. 7. The bars represent the average concentration across the mice on each day. The dots are the individual values for each mouse.

**Extended Data Fig. 9.**
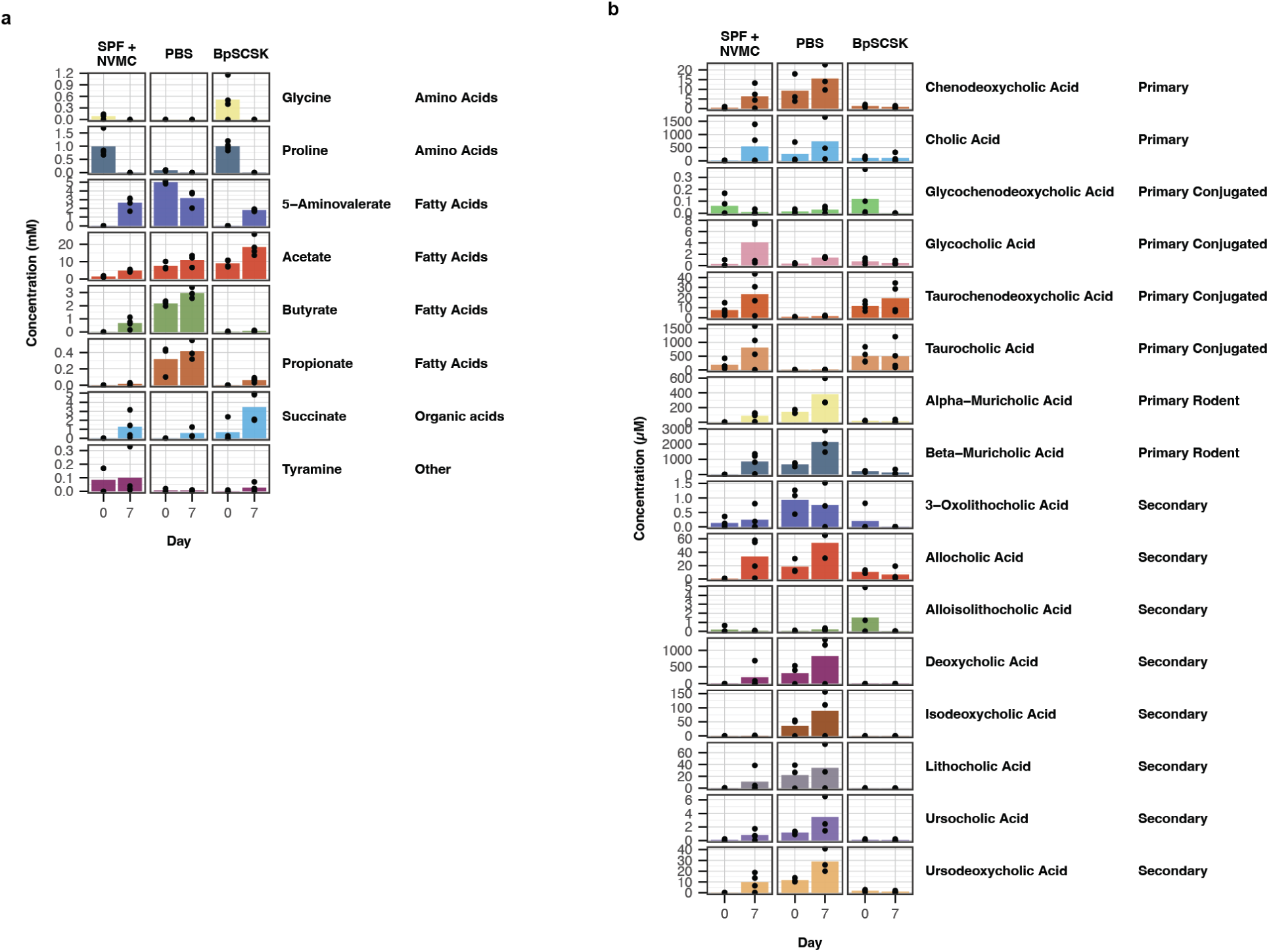
Metabolite concentrations BpSCSK-colonized mice challenged with *C. difficile*. a, b, Absolute concentrations of metabolites for mice challenged with *C. difficile* from Fig. 7. The bars represent the average concentration across the mice on each day. The dots are the individual values for each mouse.

**Extended Data Table 1.**
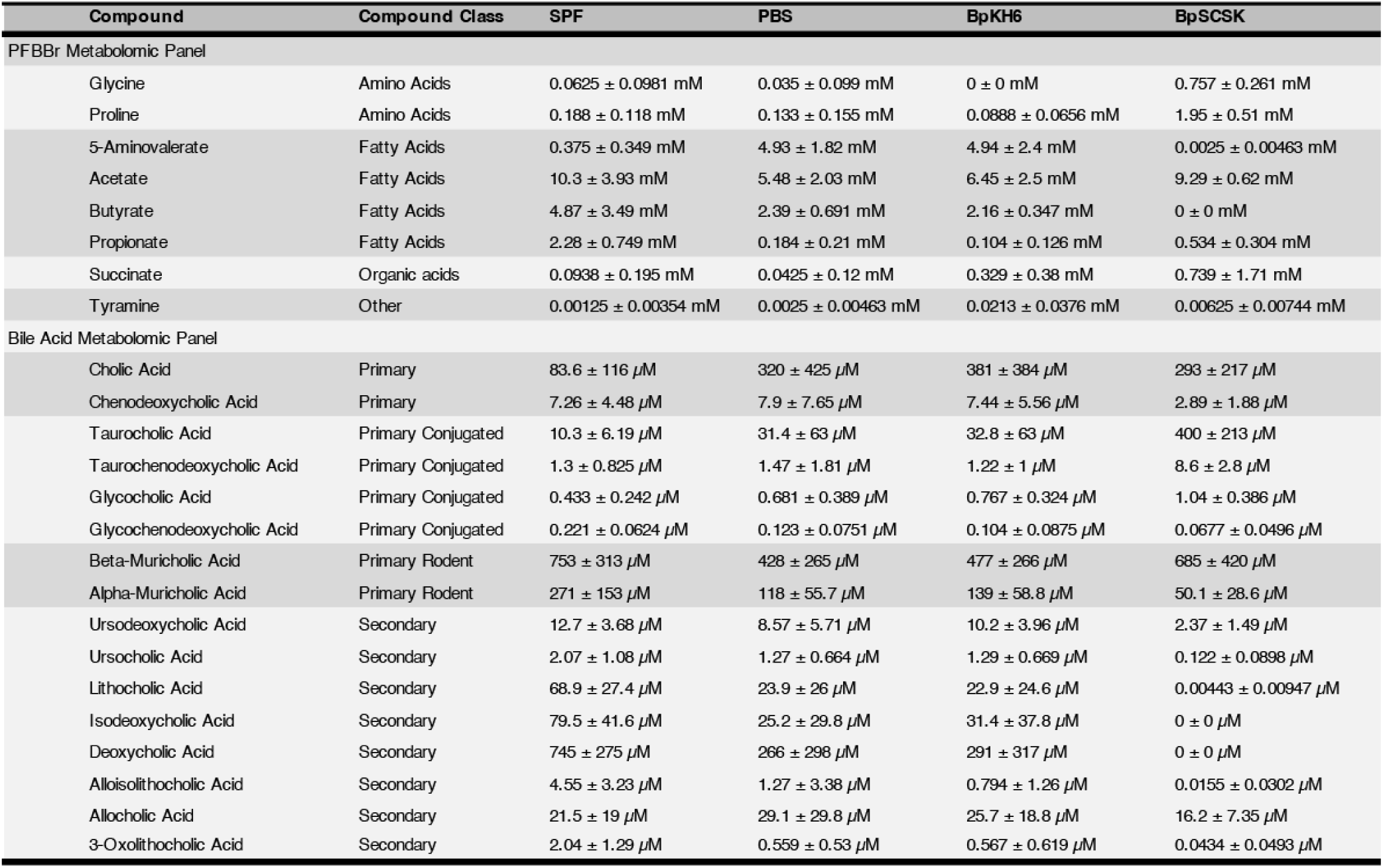
Metabolite concentrations on day 28 of colonization. Metabolite concentrations of mice from Fig. 4. Concentrations are listed as the average concentration ± standard deviation across 8 mice.

